# Epigenome profiling identifies H3K27me3 regulation of extra-cellular matrix composition in human corticogenesis

**DOI:** 10.1101/2024.10.01.616076

**Authors:** Nora Ditzer, Ezgi Senoglu, Theresa M. Schütze, Aikaterina Nikolaidi, Annika Kolodziejczyk, Katrin Sameith, Sevina Dietz, Razvan P. Derihaci, Cahit Birdir, Anne Eugster, Mike O. Karl, Andreas Dahl, Pauline Wimberger, Franziska Baenke, Claudia Peitzsch, Mareike Albert

## Abstract

Epigenetic mechanisms regulate gene expression programs during neurogenesis, but the extent of epigenetic remodelling during human cortical development remains unknown. Here, we characterize the epigenetic landscape of the human developing neocortex by leveraging Epi-CyTOF, a mass cytometry-based approach for the simultaneous single cell analysis of more than 30 epigenetic marks. We identify H3K27me3, deposited by Polycomb Repressive Complex 2 (PRC2), as the modification with the strongest cell type-specific enrichment. Inhibition of PRC2 in human cortical organoids resulted in a shift of neural progenitor cell (NPC) proliferation towards differentiation. Cell type-specific profiling of H3K27me3 not only identified neuronal differentiation genes in the human neocortex, but also extra-cellular matrix (ECM) genes. PRC2 inhibition resulted in increased production of the proteoglycan Syndecan 1. Overall, this study comprehensively characterizes the epigenetic state of specific neural cell types and highlights a novel role for H3K27me3 in regulating the ECM composition in the human developing neocortex.

## Introduction

The neocortex, responsible for higher cognitive abilities in humans, develops from a pool of NPCs that gives rise to a range of diverse cell types present in the adult brain. Cortical development involves a tight balance of NPC self-renewal and differentiation, ensuring appropriate cellular output and brain size (Borrell and Reillo, 2012; Florio and Huttner, 2014; Lui et al., 2011; Pollen et al., 2023).

The proliferative capacity of distinct NPC types differs greatly. Radial glia (RG) have the ability to self-renew, thereby replenishing the progenitor pool, and to give rise to intermediate progenitor cells (IPs) that are more neurogenic and rapidly commit to neuronal fate (Hoye and Silver, 2020; Nano and Bhaduri, 2022; Vanderhaeghen and Polleux, 2023). Underlying these cell fate transitions are precise temporal and spatial gene expression patterns that are tightly controlled by epigenetic regulation (Albert and Huttner, 2018; Amberg et al., 2019; Hirabayashi and Gotoh, 2010; Yao et al., 2016). Among diverse epigenetic mechanisms, posttranslational histone modifications occur in a huge diversity and fulfil important functions (Millan-Zambrano et al., 2022; Turner, 1993).

Especially, proteins of the Trithorax and Polycomb systems were shown to play major roles in neural stem cell differentiation (Corley and Kroll, 2015; Tsuboi et al., 2019). This group of epigenetic writers constitutes an evolutionarily conserved system, displaying antagonistic functions to maintain active and repressed chromatin states (Piunti and Shilatifard, 2016; Schuettengruber et al., 2017). Polycomb group (PcG) proteins have been implicated in the regulation of developmental timing and transcriptional programs in the mouse developing cortex (Albert et al., 2017; Amberg et al., 2022; Pereira et al., 2010; Telley et al., 2019).

Less is known about the regulation of human neocortex development, yet, many epigenetic factors have been linked to developmental disorders associated with brain phenotypes and intellectual disability (Bolicke and Albert, 2022; Mastrototaro et al., 2017). Only recently, the first studies in advanced human model systems were performed. PcG regulation was shown to control the exit from pluripotency in early human brain organoids (Zenk et al., 2024). Moreover, in concert with two other epigenetic modifiers, PcG regulation was shown to establish an epigenetic barrier in NPCs that mediates the protracted maturation of human neurons compared to other species (Ciceri et al., 2024).

This recent observation indicated that epigenetic regulation in human developing brain could be different from other species. However, currently, we lack a clear understanding of the complexity of epigenetic remodelling during human cortical neurogenesis. Such knowledge will not only reveal potential roles for epigenetic regulation in cortical evolution but may also enable future research on diagnostic and therapeutic approaches for human disorders involving epigenetic changes. Powerful single cell technologies are available for the analysis of gene expression, open chromatin, DNA methylation and 3D chromatin structure, even in combination (Noack et al., 2023), whereas single cell analysis of histone modifications is just emerging, but typically limited to few modifications (Bartosovic et al., 2021; Meers et al., 2023; Zenk et al., 2024).

Here, we apply an innovative highly multiplexed approach, Epi-CyTOF (Cheung et al., 2018; Harpaz et al., 2022), to study a large number of epigenetic modifications at single cell level in the human developing neocortex. We then focus on H3K27me3, the modification that displayed the strongest differences between neural cell types. Based on pharmacological inhibition of PRC2 and cell type-specific profiling of H3K27me3 across the genome, we propose a mechanism by which PcG proteins regulate cell fate in human cortical development.

## Results

### Epi-CyTOF resolves epigenetic marks at single cell resolution in the human developing neocortex

Epigenetic regulation of gene expression involves many different layers, including a large number of posttranslational histone modifications. To identify the histone marks that display global changes across neural cell types of the human developing neocortex, and that may therefore be important for neural cell fate, we adopted a cytometry by time-of-flight-based method for epigenetic analysis, termed Epi- CyTOF (Cheung et al., 2018; Harpaz et al., 2022) (Figure 1A). We have designed a custom panel of metal isotope-conjugated antibodies targeting 10 cell type markers, to resolve the cellular heterogeneity of the developing neocortex, and to 31 histone modifications and epigenetic enzymes (Figure 1B; Table S1). Out of the 31 antibodies targeting histone modifications, 27 were selected from previous studies, in which their specificity was validated by pharmacological inhibition, knockdown or overexpression of the responsible epigenetic modifiers (Cheung et al., 2018; Harpaz et al., 2022). The antibody panel was used to stain cell suspensions of human foetal tissue (gestation week (GW) 12–14) and human cortical organoids (Qian et al., 2020) from two independent induced pluripotent stem cell (iPSC) lines (week (W) 8). Each tissue type included 4 replicates, all of which were combined by palladium-based barcoding to reduce batch effects. Clustering of viable cell populations from human foetal tissue, based on the cell type markers SOX2, TBR2 and CTIP2, separated three main cell populations, specifically RG, IPs and neurons (N) (Figure 1C, D). Cell type clusters were further confirmed by unbiased clustering using FlowSOM (Figure S1A) and were highly consistent across replicates (Figure S1B, C). Staining intensity of the cell proliferation marker KI67, which we found to be very low in neurons, and of additional cell type markers further validated the cell populations (Figure 1E, F). Human brain organoids have become an essential tool for functional studies of human neurogenesis (Bolicke and Albert, 2022; Giandomenico and Lancaster, 2017; Pasca, 2018). To assess whether our broad panel of epigenetic marks was conserved in these *in vitro* models, we also performed Epi-CyTOF analysis on human cortical organoids (Figure 1G). This revealed that cell type clustering and staining of cell type markers (Figure 1H–K and S1A–D) was highly similar to human foetal tissue.

**Figure 1.**
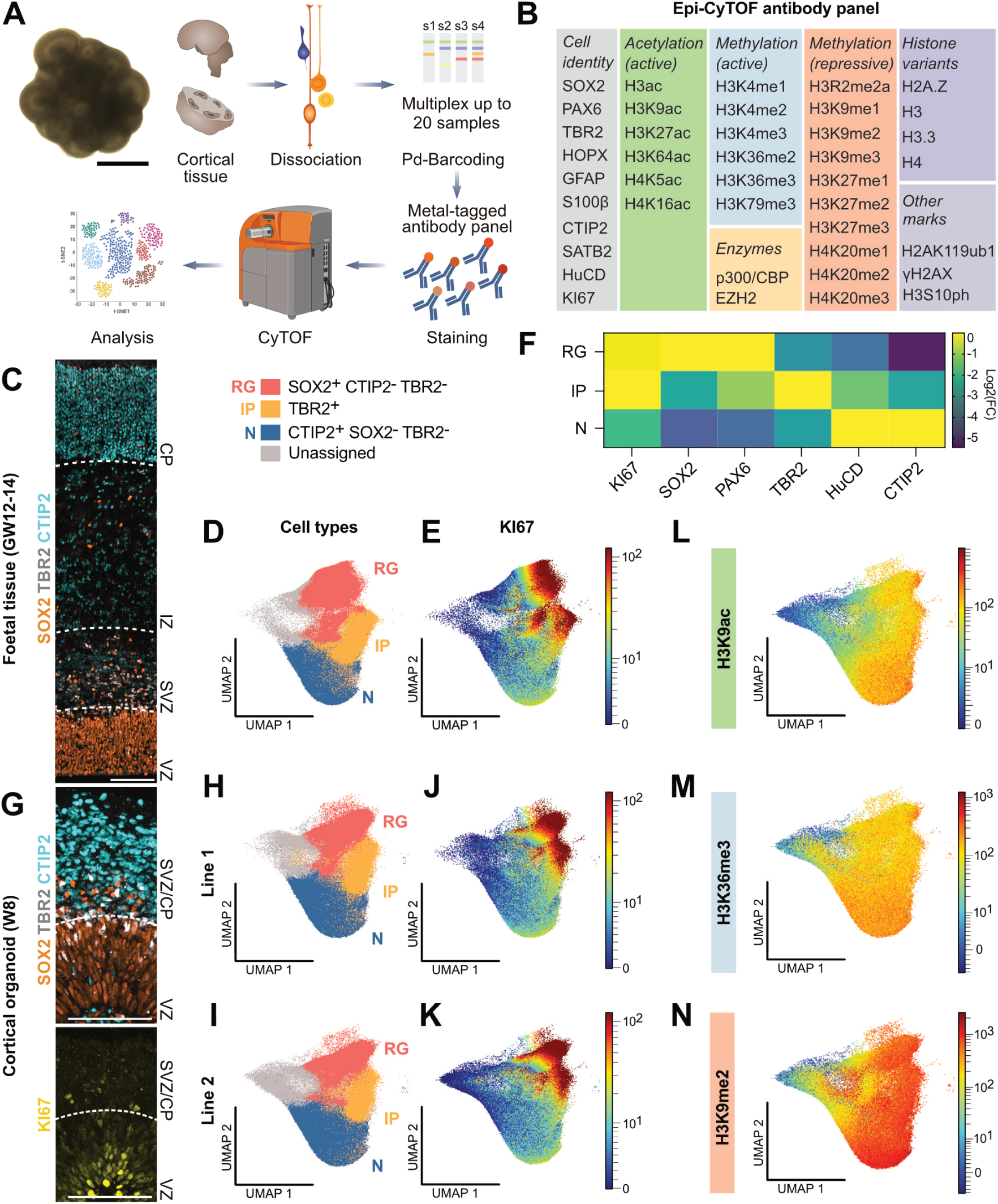
Epi-CyTOF resolves cell type-specific epigenetic profiles in the human developing neocortex. (A) Experimental workflow of epigenetic profiling by Epi-CyTOF. (B) Epi-CyTOF antibody panel for cell type and epigenetic markers. (C) Immunofluorescence staining for SOX2, TBR2 and CTIP2 of human foetal tissue (GW12–14). Scale bar, 100 µm. (D, E) UMAP analysis of cell type clustering (D) and colour-continuous scatter plots of the cell proliferation marker KI67 (E) for human foetal tissue. Scale represents metal isotope tag intensity. (F) Heat map of Log2 fold changes of median metal tag intensities for cell type markers in RG, IPs and neurons (N) from foetal tissue (4 samples from 2 different individuals). (G) Immunofluorescence for SOX2, TBR2 and CTIP2 (top) and KI67 (bottom) of cortical organoids (W8). Scale bars, 100 µm. (H–K) UMAP of cell type clusters (H, I) and KI67 intensity (J, K) for cortical organoids (W8) generated from iPSC lines 1 and 2 (each from a combination of 4 replicates from independent organoid batches). (L–N) UMAP colour-continuous scatter plots for selected markers of active acetylation (L; H3K9ac), active methylation (M; H3K36me3) and repressive methylation (N; H3K9me2) marks.

Next, we analysed Epi-CyTOF data of histone modifications in human foetal tissue (Figure 1L–N) and cortical organoids (Figure S1E), resulting in single cell epigenetic information. The staining patterns for active histone acetylation (H3K9ac), and active (H3K36me3) and repressive (H3K9me2) histone methylation marks were consistent between primary tissue and *in vitro* models, but differed between neural cell types, displaying higher enrichment in neurons compared to RG and IPs.

We then determined the mean metal tag intensities for all 31 epigenetic readouts for RG, IPs and neurons for each individual replicate (Figure S2A–C), highlighting the power of the methodology to determine changes in global levels of a large number of epigenetic modifications across 12 samples at once. This revealed that many modifications display differential enrichment across different neural cell types.

Taken together, Epi-CyTOF is not only able to resolve the cellular heterogeneity of the human developing neocortex, but also provides single cell epigenetic information for a large number of posttranslational histone modifications.

### H3K27me3 is highly enriched in cortical neurons compared to NPCs

To identify the posttranslational histone modifications that most strongly change from RG to neurons, we plotted the fold change of metal tag intensities for all epigenetic markers analysed by Epi-CyTOF (Figure 2A and S2D–H). In human foetal tissue, 21 of the 31 markers had a higher staining intensity in neurons compared to RG, including both active and repressive histone modifications (Figure 2A). The remaining modifications displayed various patterns of enrichment, with several modifications showing the highest levels in IPs (Figure S2F–H). Among all histone markers, H3ac was the only modification that showed higher metal tag intensity in RG compared to both IPs and neurons in human foetal tissue. This is in line with the view that epigenetic marks accumulate with progression of cell fate commitment during lineage specification (Barth and Imhof, 2010), and our data suggest that this involves both active and repressive histone modifications during neural differentiation in the human developing neocortex.

**Figure 2.**
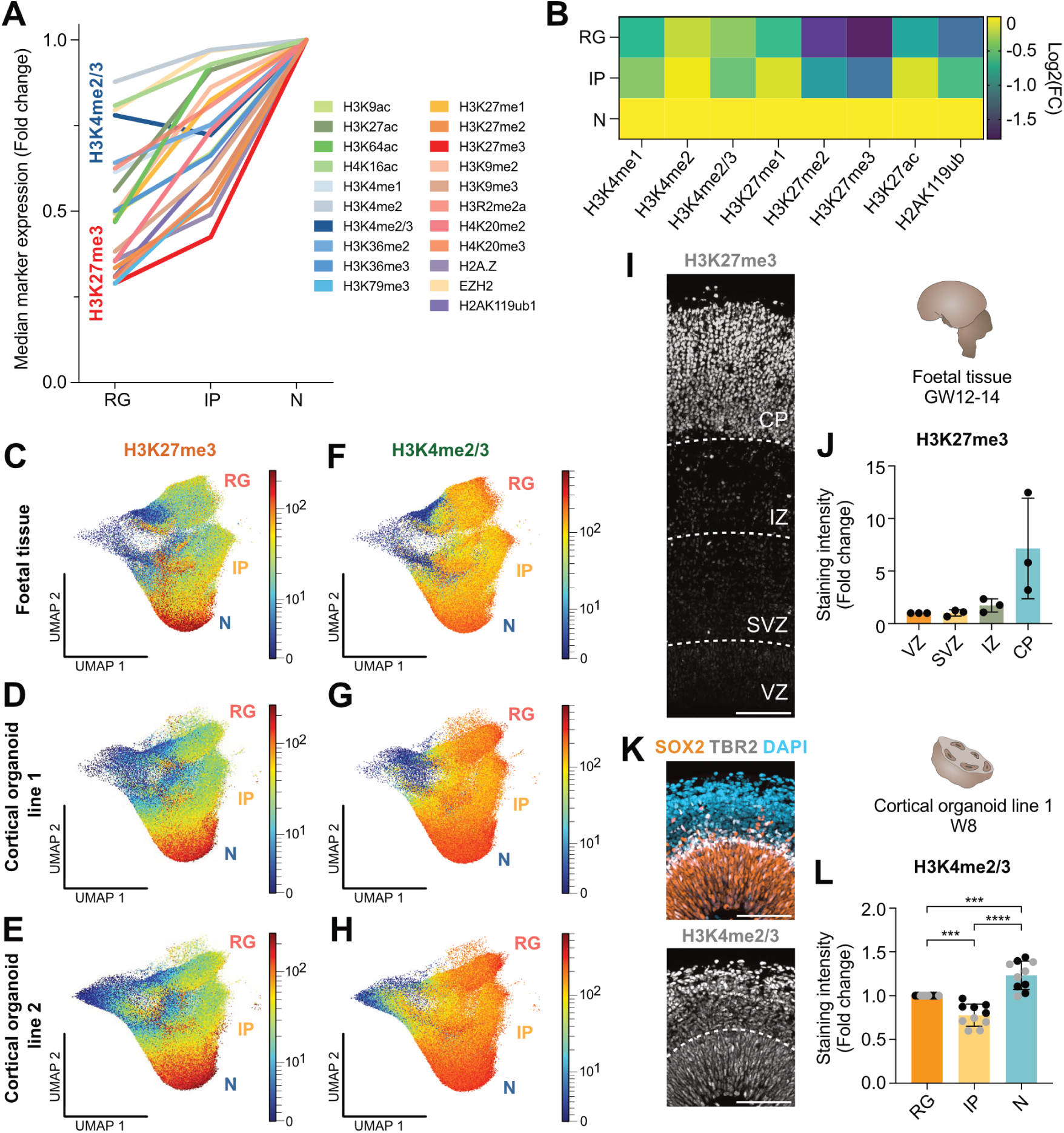
Epi-CyTOF reveals a strong enrichment of H3K27me3 in human cortical neurons compared to NPCs. (A) Line plot displaying epigenetic marks with increasing abundance from RG to N. Values represent fold change of Epi- CyTOF median metal tag intensities in foetal tissue (GW12-14). (B) Heat map of Log2 fold changes of median metal tag intensities for epigenetic marks in RG, IP and N from foetal tissue (4 samples from 2 different individuals). (C–H) UMAP colour-continuous scatter plots for the intensity of H3K27me3 (C–E) and H3K4me2/3 (F–H) in foetal tissue (C, F) and of cortical organoids from iPSC line 1 (D, G) and 2 (E, H) (W8). (I) Immunofluorescence of H3K27me3 of human foetal tissue (GW12–14). (J) Quantification of H3K27me3 intensity in human foetal tissue in the ventricular zone (VZ), sub-ventricular zone (SVZ), intermediate zone (IZ), and cortical plate (CP) relative to the VZ. (K) Immunofluorescence for SOX2, TBR2 and CTIP2 (top) and H3K4me2/3 (bottom) of cortical organoids (W8). (L) Quantification of H3K4me2/3 intensity in RG, IPs and N in cortical organoids from iPSC line 1 relative to RG. Scale bars, 100 µm. Bar graphs represent mean values. Error bars represent SD; J, of 3 tissue samples from independent individuals; L, of 10 organoids from 2 batches.

The histone modification that showed the strongest change from RG to neurons in human foetal tissue was H3K27me3 (Figure 2A). This modification also strongly changed from RG to neurons in cortical organoids from both iPSC lines (Figure S2D, E). H3K27me3 is a repressive modification mediated by PRC2 and counteracted by H3K4me3, which is associated with active transcription (Piunti and Shilatifard, 2016; Schuettengruber et al., 2017). Additional Polycomb-associated modifications, specifically H3K27me2 and H2AK119ub, also displayed high differential enrichment in neurons compared to RG (Figure 2A, B). In contrast, H3K4me2/3 was detected in all cell types, with a slight decrease from RG to IPs, accompanied by a minor increase in neurons. The patterns of H3K27me3 and H3K4me2/3 staining intensity across the three major cell type clusters were highly similar in human foetal tissue and cortical organoids from both iPSC lines (Figure 2C–H).

To independently verify these findings, we performed immunohistochemistry for H3K27me3 on human foetal tissue, using a different primary antibody (Figure 2I). Quantification of the staining intensity confirmed the low levels of H3K27me3 in the ventricular (VZ) and subventricular (SVZ) zone, containing RG and IPs, respectively, and the strong enrichment of H3K27me3 in the cortical plate (CP), where neurons reside (Figure 2I).

Likewise, immunohistochemistry for H3K4me2/3 of human cortical organoids confirmed the Epi- CyTOF staining pattern, with slightly lower levels in IPs compared to RG and neurons (Figure 2K, L), indicating that Epi-CyTOF is able to detect small differences in histone modifications between cell types. Overall, these findings suggest that analysis of histone modifications by Epi-CyTOF was highly robust across multiple replicates in complex tissue samples and could be independently confirmed for selected epigenetic marks in human foetal tissue and cortical organoids.

To further investigate the differential enrichment pattern of H3K27me3 in the human developing neocortex, we asked whether PRC2 components were also differentially expressed. Both gene expression analysis of published transcriptome datasets (Florio et al., 2015) and immunohistochemistry for EZH2 and SUZ12 indicated that PRC2 core components were uniformly expressed across the germinal zones and cortical plate of the human foetal neocortex (Figure S3A–C). Immunohistochemistry across a time course of human cortical organoid development showed that PRC2 expression was also largely uniform in the VZ compared to the CP containing neurons in organoids (Figure S3D-F). Interestingly, however, while H3K27me3 is initially homogenously distributed throughout the ventricle-like structures of human cortical organoids (W2–4), H3K27me3 is strongly enriched in neurons compared to progenitors from W6 onwards, despite uniform distribution of PRC2 (Figure S3D-F), mirroring the pattern in human foetal tissue (Figure 2I, J).

In summary, Epi-CyTOF identified PRC2-mediated H3K27me3 as the most differentially distributed histone modification between RG and neurons, which we independently confirmed in both human foetal tissue and cortical organoids. During cortical organoid development, H3K27me3 was initially uniformly distributed throughout the ventricle-like structures. However, from early stages of neurogenesis it appeared strongly enriched in neurons compared to NPCs.

### Genome-wide H3K27me3 profiles distinguish neural cell populations

We next aimed to elucidate the distribution of H3K27me3 and H3K4me2/3 throughout the genome in specific cell types of the human developing neocortex. Therefore, we isolated nuclei from human foetal tissue (GW12–14), sorted the three major cell populations (RG, IPs and neurons) using fluorescence activated nuclei sorting (FANS) and performed Cleavage Under Targets and Tagmentation (CUT&Tag) (Kaya-Okur et al., 2019) (Figure 3A). RG, IPs and neurons were sorted based on expression of the cell type markers PAX6, TBR2 and CTIP2, respectively (Figure S4A–B). Cell identity of the sorted populations was further validated by expression analysis of additional marker genes (Figure S4C). Moreover, to prevent potential unspecific signal from the sorting antibodies in CUT&Tag, we applied a blocking step using Fab-fragments (Ahanger et al., 2024). Successful blocking was confirmed by the lack of regions significantly enriched for H3K27me3 in an unstained versus stained cell population (Figure S4D).

**Figure 3.**
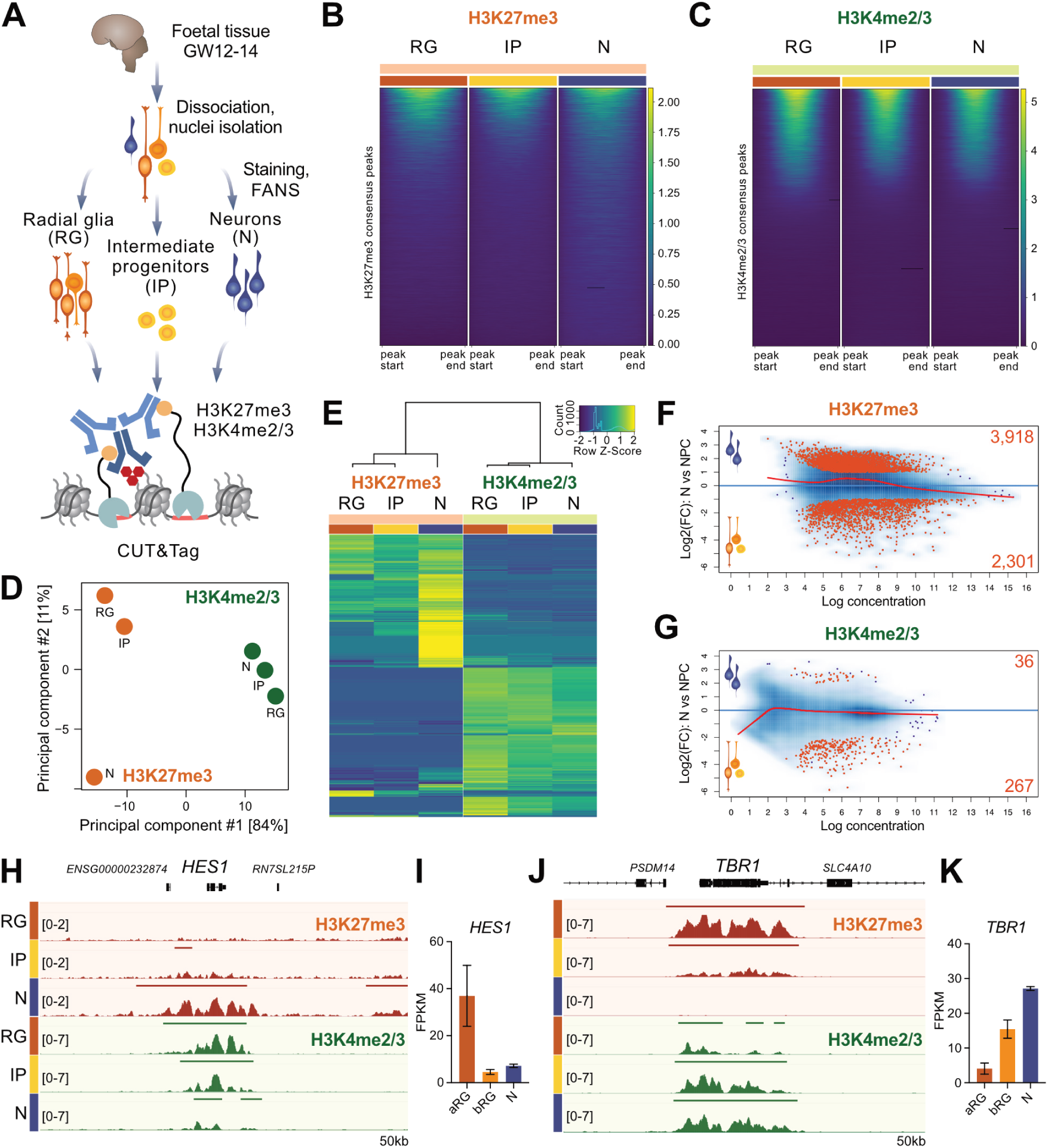
H3K27me3 profiles distinguish NPCs and neurons in the human developing neocortex. (A) Experimental scheme for cell type-specific profiling of histone marks by CUT&Tag acquired for RG, IP and N from human foetal tissue (GW12–14; two tissue samples from independent individuals). (B, C) Heatmaps of the enrichment of H3K27me3 (B) and H3K4me2/3 (C) acquired for RG, IP and N by CUT&Tag in a set of consensus peaks. Each column represents the mean of two samples. (D) PCA analysis for H3K27me3 and H3K4me2/3. Each dot represents the average of two samples. (E) Heatmaps for H3K27me3 or H3K4me2/3 peaks. Plotted are Z-scores for the top 20K regions, sorted by standard deviation. Each column represents the average of two samples. (F, G) MA plot illustrating differential enrichment of H3K27me3 (F) and H3K4me2/3 (G) between N and NPCs (combining RG and IP), determined by DESeq2. Regions significantly enriched for one of the histone marks in one cell population are marked in red (p-adjust < 0.05, logFC > 1).(H) Tracks of H3K27me3 and H3K4me2/3 from RG, IP and N at the *HES1* genomic locus. (I) Corresponding *HES1* mRNA levels in apical radial glia (aRG), basal radial glia (bRG) and N from human foetal cortex analysed by RNA-seq (data from (Florio et al., 2015)). (J) Tracks of H3K27me3 and H3K4me2/3 at the neuronal gene locus *TBR1*. (K) *TBR1* mRNA levels analysed by RNA-seq.

Applying this cell type-specific CUT&Tag strategy, we found H3K27me3, but not H3K4me2/3, to display a distinct genome wide distribution in neurons compared to RG and IPs (Figure 3B, C). Whereas a small set of regions was strongly enriched for H3K27me3 in RG and IPs, H3K27me3 was enriched at low levels at additional regions in neurons (Figure 3B). In contrast, the distribution pattern of H3K4me2/3 was more similar between cell types (Figure 3C). This was also reflected in the principal component analysis (PCA), in which H3K27me3 of RG and IPs was clearly separated from neurons, whereas all three cell types clustered together for H3K4me2/3 (Figure 3D). Z-scores of the top 20,000 regions for H3K27me3 or H3K4me2/3 further confirmed this observation (Figure 3E).

To identify differentially enriched regions for H3K27me3 and H3K4me2/3 in individual cell types, we used DESeq2 (p-adjust ≤ 0.05; logFC ≥ 1). Since we did not detect any regions specifically enriched for H3K27me3 in IPs, we pooled RG and IPs, referred to as ‘NPCs’. In total, 3,918 neuron- and 2,301 NPC-specific regions were detected for H3K27me3 (Figure 3F; Table S2). Compared to the NPC-specific regions, neuron-specific regions displayed, on average, lower enrichment of H3K27me3. In contrast, only 36 neuron- and 267 NPC-specific regions were detected for H3K4me2/3 (Figure 3G) despite the overall higher number of binding sites marked by H3K4me2/3 compared to H3K27me3 (Figure S4E, F).

Cell type-specific distribution of H3K27me3 was, for example, found at the transcription factor gene *HES1* that became silenced throughout neuronal differentiation and concomitantly acquired low levels of H3K27me3 (Figure 3H, I). In contrast, the neuronal gene *TBR1* showed high enrichment of H3K27me3 in RG, which was depleted upon neuronal fate commitment (Figure 3J, K).

Overall, these results are in line with the global levels of H3K27me3 and H3K4me2/3 characterized by Epi-CyTOF. Using cell type-specific epigenomics, we found the repressive H3K27me3 mark to become progressively enriched throughout the neuronal differentiation trajectory, while we did not observe comparably dynamic changes for the active H3K4me2/3 mark. The H3K27me3 profile clearly distinguished NPCs and neurons, with neurons depicting a higher number of PRC2 target genes. Interestingly, neuron-specific H3K27me3 binding sites carried lower enrichment of the mark compared to high enrichment of H3K27me3 at NPC-specific sites.

### Inhibition of PRC2 induces a shift in NPC proliferation towards differentiation in human cortical organoids

Both, the global enrichment of H3K27me3 in neurons as well as the cell type-specific genome-wide distribution of H3K27me3, suggest a role of PRC2 in regulating cell fate transitions in the human developing neocortex. Prior to this study, the role of PRC2 in regulating neuronal maturation has been described in detail (Ciceri and Studer, 2024). Thus, we focussed on functionally investigating the role of PRC2 in NPCs throughout human corticogenesis. We applied pharmacological inhibition of EED (Qi et al., 2017), a core subunit of PRC2, to cortical organoids from W3 to W8 of culture (Figure 4A). The treatment was performed using 5 µM EED226 (EEDi), as this was the concentration that resulted in an almost complete depletion of H3K27me3 compared to the DMSO control after 5 weeks of treatment, without compromising organoid development (Figure 4B, C and S5A). H3K27me3 was also abolished in neurons, which have a very high enrichment of this histone methyl mark to begin with and represent post-mitotic cells, that therefore do not dilute histone modifications during DNA replication.

**Figure 4.**
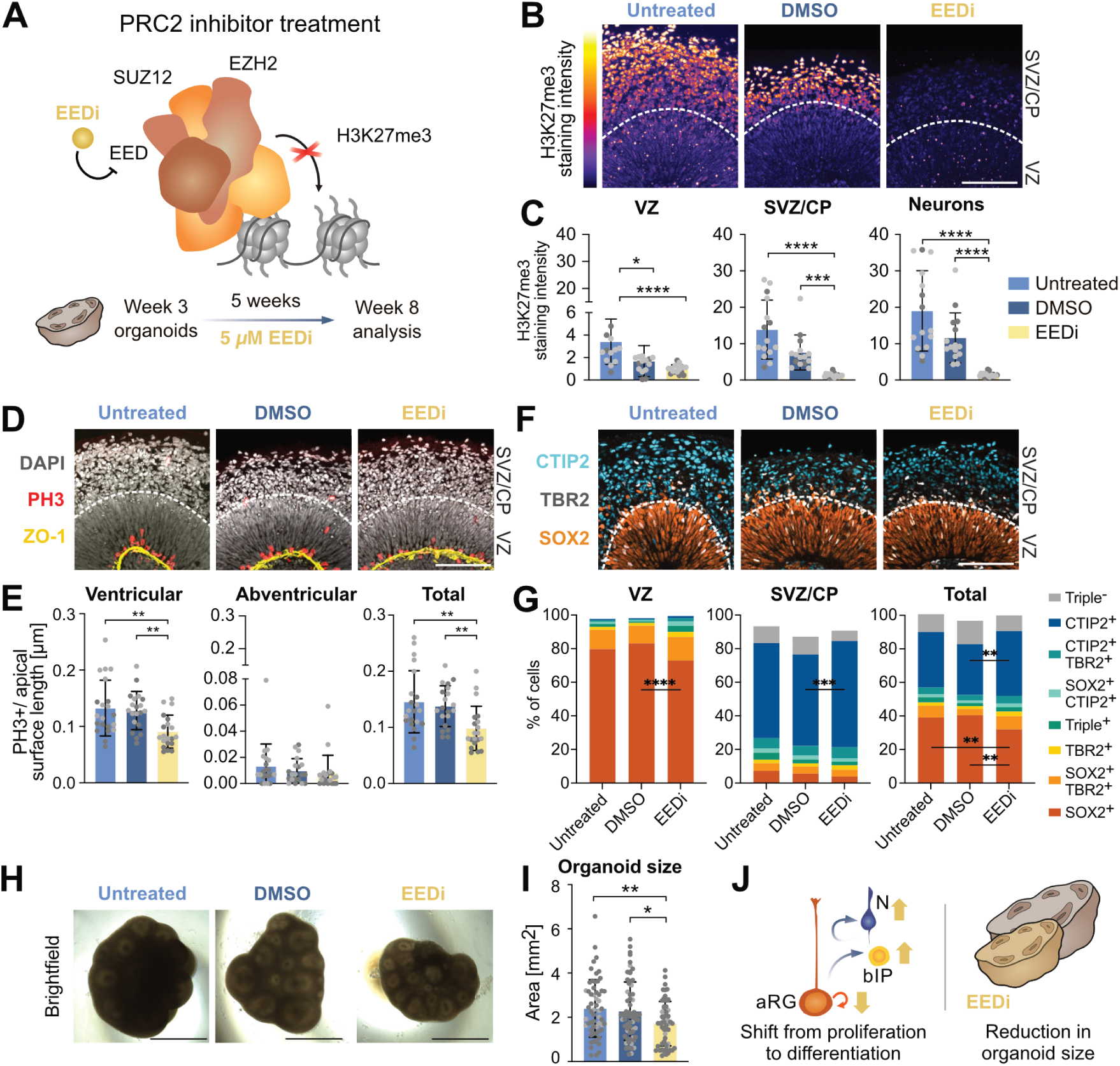
Inhibition of PRC2 results in a shift of NPC proliferation to differentiation in the human developing neocortex. (A) Experimental scheme for pharmacological inhibition of PRC2 in human cortical organoids. Treatment was performed with 5 µM EED inhibitor EED226 (EEDi) from week 3 to week 8 of organoid culture. No treatment (‘Untreated’) and DMSO treatment (‘DMSO’) serve as controls. (B) Immunofluorescence of H3K27me3 in cortical organoids treated with EEDi (W8). (C) Quantification of H3K27me3 staining intensity in nuclei (segmented using DAPI) of VZ, SVZ/CP and CTIP2-positive N. (D) DAPI staining and immunofluorescence of the apical surface marker ZO-1 and PH3. (E) Quantification of PH3+ cells per apical surface length (determined by ZO-1 signal) in ventricular (up to three nuclei away from the apical surface), abventricular and total area. (F) Immunofluorescence of SOX2, TBR2 and CTIP2. (G) Quantification of SOX2, TBR2 and CTIP2 as percentage of total cells (determined by DAPI) in VZ, SVZ/CP and total area. (H) Brightfield images of cortical organoids treated with EEDi (W8). (I) Quantification of total organoid size based on bright field images (as in H). (J) Summary scheme of phenotypes induced by pharmacological inhibition of PRC2. Scale bars, (B, D, F) 100 µm, (H) 1 mm. Bar graphs represent mean values. Error bars represent SD; C, of 17–19 ventricles from at least 10 organoids; D, of 18–20 ventricles from at least 10 organoids.; G, of 12 ventricles from at least 8 organoids; I, of 56–63 organoids from 2 organoid batches (indicated by different colours). **** p < 0.0001, *** p < 0.001, ** p < 0.01, * p < 0.05; Kruskal-Wallis test with Dunn’s post hoc test (C, E), two-way (G) or one-way (I) ANOVA with Tukey’s post-hoc test.

To assess the effects of H3K27me3 depletion on cortical organoid development, we first analysed the mitotic marker phospho-histone 3 (PH3) and found a significant decrease in ventricular and total mitosis compared to the untreated and DMSO controls (Figure 4D, E). As this could indicate a reduced potential of NPCs for proliferation, we next examined the cell proliferation marker KI67. While we did not observe a reduction in the total percentage of KI67-positive cells, the percentage of KI67-positive cells was reduced in the SVZ/CP (Figure S5B, C), where IPs reside. This was mirrored by an increase in the percentage of PAX6/TBR2 double-positive cells among total KI67^+^ cells, at the cost of PAX6 single-positive cells (Figure S5D, E).

To test if this shift towards a higher percentage of neurogenic IPs resulted in increased neuronal differentiation, we performed immunohistochemical analysis for the RG marker SOX2, the IP marker TBR2 and the neuronal marker CTIP2 (Figure 4F, G). We found that loss of H3K27me3 resulted in a significant reduction of SOX2 single-positive RG in the VZ. Moreover, consistent with previous results of EZH2 inhibition in human forebrain organoids (Ciceri et al., 2024), we observed a significant increase in CTIP2-positive neurons in the SVZ/CP of EEDi-treated cortical organoids compared to the DMSO control.

So far, these observations suggest that loss of H3K27me3 induces a shift of NPC proliferation towards differentiation in human cortical organoids. Such a shift would eventually be expected to result in a lower neuronal output due to a reduction of the NPC pool. Indeed, total organoid size was significantly reduced by EEDi treatment (Figure 4H, I), accompanied by a reduction in the VZ thickness in ventricle-like regions (Figure S5F, G). Staining for cleaved caspase 3 (CC3) confirmed that this change in size was not caused by increased apoptosis in the inhibitor treated organoids (Figure S5H, I). Taken together, these data reveal a role for PRC2 in maintaining human NPCs in a proliferative state throughout cortical development, thus ensuring appropriate neuronal output and cortical organoid size (Figure 4J).

### H3K27me3 is enriched on neuronal differentiation and ECM genes

Identification of PRC2 as a regulator of human NPC cell fate decisions led us to ask which genes were modified by H3K27me3 and how these targets might mediate the phenotypes observed upon PRC2 inhibition. Therefore, we performed a detailed analysis of cell type-specific PRC2 targets in RG, IPs and neurons isolated from the human foetal neocortex (Figure 5A). Overall, a higher number of H3K27me3 binding sites were found in neurons compared to RG and IPs (Figure 5B). This was reflected in a higher number of significantly enriched neuron-specific target regions, identified by differential analysis of H3K27me3 (Figure 5C).

**Figure 5.**
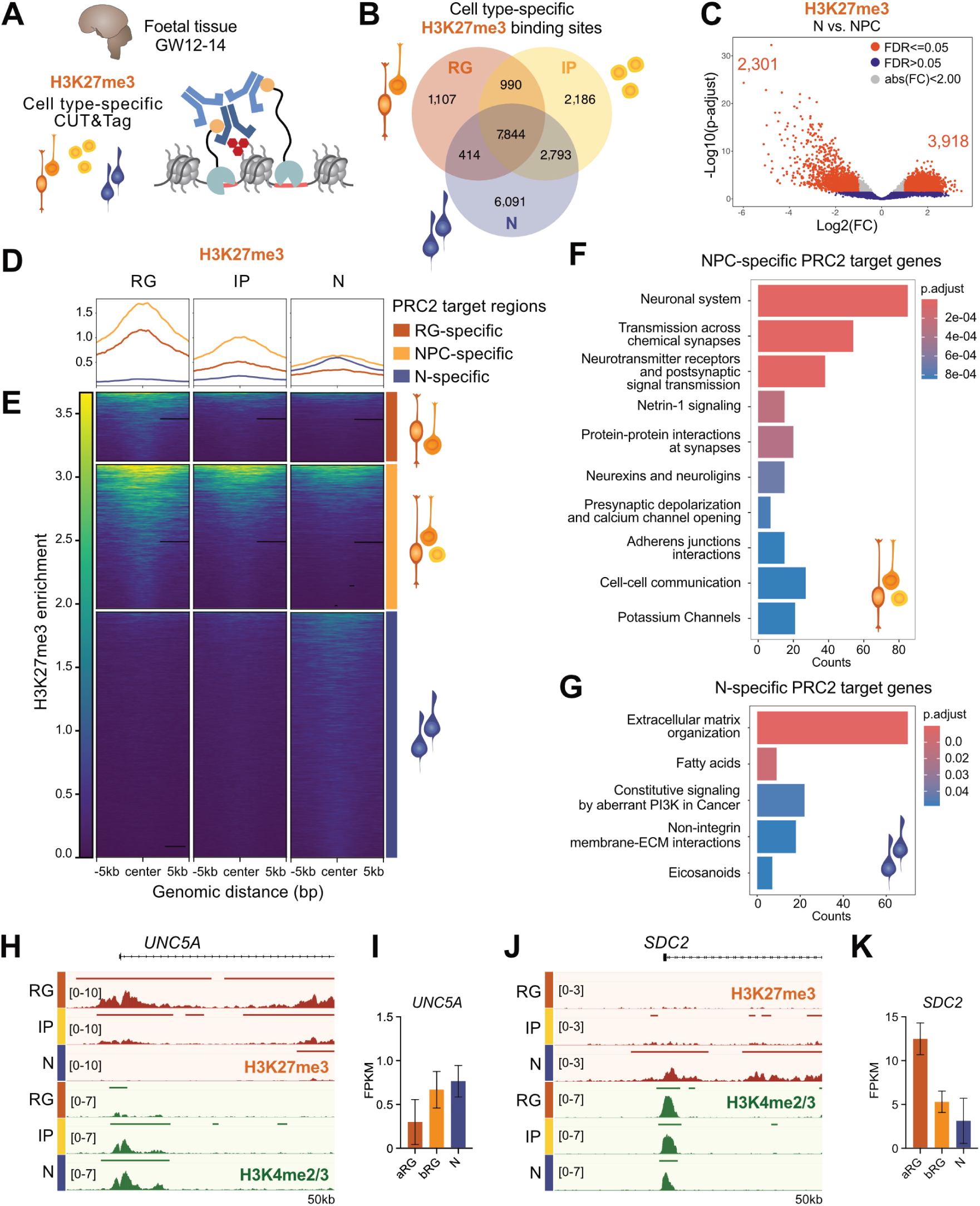
H3K27me3 is enriched on neuronal differentiation and ECM genes. (A) Experimental scheme for cell type-specific profiling of histone marks by CUT&Tag acquired for RG, IP and N from human foetal tissue (GW12–14; two tissue samples from independent individuals). (B) Venn diagram showing the overlap of H3K27me3 binding sites in RG, IP and N. (C) Volcano plot displaying regions of differential H3K27me3 enrichment between N and NPC (combining RG and IP). Regions significantly enriched in one cell population are marked in red (p-adjust < 0.05, logFC > 1). (D, E) Profiles (D) and heatmaps (E) of the H3K27me3 enrichment of RG-, NPC- and N-specific PRC2 target regions, determined by DESeq2. Each column represents the average of two samples. (F, G) Reactome pathway enrichment for NPC- (E) and N-specific (F) PRC2 target genes. (H) Tracks of H3K27me3 and H3K4me2/3 from RG, IP and N at the *UNC5A* genomic locus. (I) Corresponding *UNC5A* mRNA in human foetal cortex analysed by RNA-seq (data from (Florio et al., 2015)). (J) Tracks of H3K27me3 and H3K4me2/3 from RG, IP, and N at the *SDC2* genomic locus. (K) *SDC2* mRNA levels analysed by RNA-seq.

Plotting of the average coverage of H3K27me3 revealed that neuron-specific target regions, while more abundant, displayed lower enrichment of the repressive histone mark compared to NPC-specific targets (Figure 5D, E). NPC-specific target regions showed a strong depletion of H3K27me3 upon differentiation. In contrast, neuron-specific targets acquired comparably lower levels of the repressive mark, once neuronal fate was established (Figure 5D, E).

To examine the function of the cell type-specific PRC2 targets, we annotated each region to its nearest neighbouring gene and performed pathway analysis (Yu and He, 2016). As expected, in NPCs, PRC2 was found to regulate genes associated with neuronal differentiation processes (Figure 5F). In addition, pathway analysis pointed towards secreted signalling molecules, including netrins and their receptors. Moreover, in neurons, PRC2 target genes were linked to ECM organization and ECM interactions (Figure 5G). These findings are especially interesting considering previous results from mice, where cortex-specific knockout of *Eed* led to changes in the neurogenic differentiation of NPCs only upon tissue-wide knockout, but not upon deletion of *Eed* in individual cells in an otherwise wildtype tissue (Amberg et al., 2022). Therefore, these results suggest a non-cell autonomous mechanism for the regulation of NPC fate decisions by PRC2.

Among the genes regulated by PRC2 in a cell type-specific manner and encoding signalling molecules or ECM components was *Unc-5 netrin receptor A* (*UNC5A*) (Figure 5H, I). This NPC-specific target gene was repressed by H3K27me3 in RG and IPs, and became activated in neurons. Netrin signalling has been described to direct cell migration and axon migration in the developing brain (Moore et al., 2007), thus actively regulating tissue morphology and structure. Among the neuron-specific PRC2 target genes, we identified different syndecans, which are cell surface proteoglycans that have been linked to the regulation of neural stem cell proliferation in various studies (Mouthon et al., 2020; Wang et al., 2012). Specifically, *Syndecan 2* (*SDC2*) was expressed in RG, but was downregulated and enriched in H3K27me3 in neurons (Figure 5J, K).

Taken together, we identified PRC2 as a regulator of ECM composition and secretion in the human developing neocortex. Intriguingly, rather than only targeting neuronal differentiation genes directly and preventing differentiation in a cell-autonomous manner, PRC2 might have a secondary mechanism of action. By regulating the expression of ECM genes, PRC2 may contribute to the localized enrichment of pro-proliferative ECM components in the germinal zones, thus potentially maintaining NPC proliferative capacity through a non-cell autonomous process.

### PRC2 inhibition results in aberrant ECM production in cortical organoids

To evaluate the effect of PRC2 inhibition on histone methylation at target genes, we performed cell type-specific CUT&Tag for H3K27me3 and H3K4me2/3 of EEDi treated organoids (Figure 6A). This confirmed our results obtained by immunohistochemistry analysis (Figure 4B, C), revealing an essentially complete loss of H3K27me3 at previously enriched regions in RG, IPs and neurons (Figure 6B). As a result, all three cell types were found clustered together in PCA analysis of H3K27me3 after EEDi treatment of organoids, whereas for DMSO control samples, RG and IPs were closer together, but separate from neurons (Figure 6C), as also previously observed for human foetal tissue (Figure 3D). Overall, we saw a very strong depletion of H3K27me3 in the EEDi versus DMSO control condition (Figure 6D), with 28,241 regions showing a significant loss of H3K27me3 (Figure 6E). Of note, loss of H3K27me3 had no significant effect on H3K4me2/3 profiles (Figure S6A–C).

**Figure 6.**
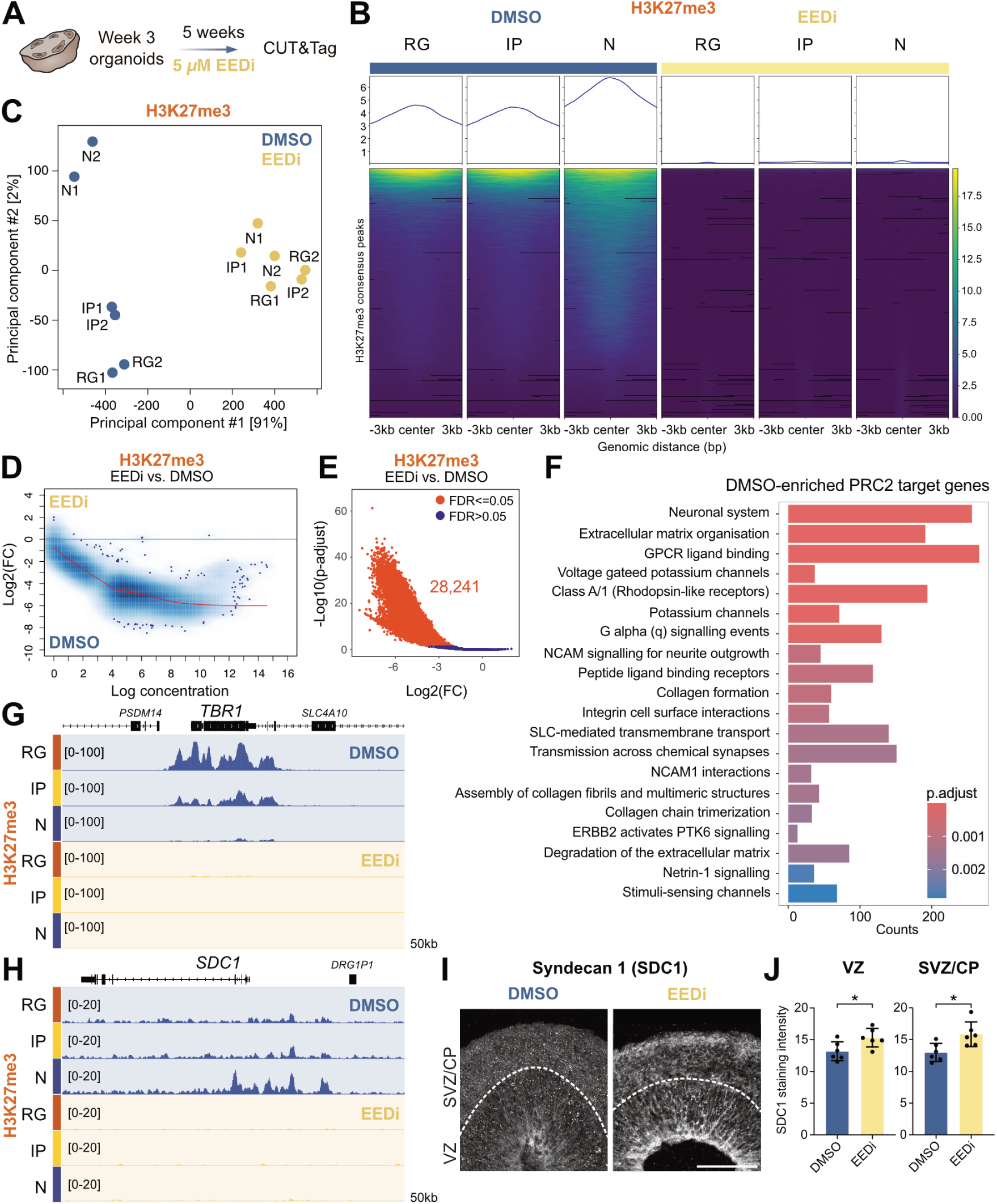
PRC2 inhibition results in aberrant ECM production in human cortical organoids. (A) Experimental scheme for pharmacological inhibition of PRC2 in human cortical organoids. (B) Profiles and heatmaps of the enrichment of H3K27me3 in RG, IP and N sorted from cortical organoids treated with EEDi or DMSO determined by CUT&Tag in a set of consensus peaks. Each column represents the mean of two independent organoid batches. (C) PCA analysis for cell type-specific H3K27me3 acquired for RG, IP and N. Each dot presents a cell type from an independent organoid batch treated with EEDi or DMSO. (D) MA plot illustrating differential enrichment of H3K27me3 between EEDi and DMSO control condition, determined by DESeq2. Significant regions are not highlighted. (E) Volcano plot displaying regions of differential H3K27me3 enrichment between EEDi and DMSO control condition. Regions significantly enriched in one condition are marked in red (p-adjust < 0.05, logFC > 1). (F) Reactome pathway enrichment for DMSO-enriched PRC2 target genes. (G, H) Tracks of H3K27me3 from RG, IP and N sorted from cortical organoids treated with EEDi or DMSO at *TBR1* (G) and *SDC1* (H) genomic loci. (I) Immunofluorescence for SDC1 in cortical organoids from DMSO and EEDi treatment conditions. (J) Quantification of SDC1 staining intensity in VZ or SVZ/CP of cortical organoids from DMSO and EEDi treatment conditions. Scale bar, 100 µm. Bar graphs represent mean values. Error bars represent SD; J, of 6 ventricles from at least 3 organoids. * p < 0.05; two-tailed unpaired t-test.

Next, we associated the regions, that were sensitive to H3K27me3 loss, to their predicted target genes. Despite the majority of H3K27me3 being lost, the pathways that were most strongly enriched were associated to neuronal systems and ECM organisation (Figure 6F). Interestingly, typical terms associated with Polycomb regulation, such as genes related to the development of other organs (heart, digestive tract, skeletal system) (Albert et al., 2017; Zenk et al., 2024), were not enriched. Among the neuronal genes that lost H3K27me3 upon EEDi treatment, was the T-Box Brain Transcription Factor *TBR1* (Figure 6G), that regulates multiple aspects of cortical development in humans, including neuronal migration, areal identity and axon projection (den Hoed et al., 2018; Deriziotis et al., 2014). Among the terms related to the extracellular environment, in addition to ECM organization, collagen formation, integrin cell surface interactions, ECM degradation and Netrin-1 signalling are noteworthy. All these terms point to a role of PRC2 target genes in remodelling the extracellular niche during human cortical development. One example is *Syndecan 1* (*SDC1*), that displayed the highest enrichment of H3K27me3 in neurons, which was lost upon EEDi treatment (Figure 6H).

To investigate the effects of H3K27me3 depletion on gene expression, we performed RNA-seq for FANS-isolated RG, IPs and neurons from the DMSO and EEDi conditions after five weeks of treatment of human cortical organoids. Overall, the effects on gene expression were minor, and after EEDi treatment, the RG, IP and neuron cell populations could still be distinguished and clustered with the DMSO control samples (Figure S6D). In line with the repressive function of PRC2, we did not detect any significantly downregulated genes, but 18 genes upregulated in all cell types (Figure S6E, refer to Table S3 for genes upregulated in individual cell types), all of which were associated with synapse function (Figure S6F). The small effect on gene expression is in agreement with previous studies in both mouse and human, identifying early embryonic development the most critical period of PRC2 function (O’Carroll et al., 2001; Pasini et al., 2004), especially during the exit from pluripotency (Boyer et al., 2006; Pasini et al., 2007; Zenk et al., 2024). Moreover, these results reflect our finding that human cortical organoids continue to develop grossly normal once neural fate is established at week 3, resulting in only small changes in cell populations at week 8 (Figure 4). The biggest difference that we observed, was the shift of NPC fate from proliferation to differentiation, which is highly dependent on the extracellular niche of the human developing neocortex (Cubillos et al., 2024; Massimo and Long, 2021).

Given the strong enrichment of ECM-related genes among PRC2 targets, we specifically asked whether this group of genes was affected upon H3K27me3 depletion. Gene set enrichment analysis (GSEA) revealed a small, not significant enrichment of ECM genes among EEDi-upregulated genes in neurons compared to RG (Figure S6G). We therefore continued to examine differences in ECM-related factors on protein level. Specifically, we stained human cortical organoid sections from the DMSO and EEDi treatment conditions for the heparan sulfate proteoglycan Syndecan 1, that we found to be cell type-specifically regulated by PRC2 (Figure 6H). Syndecan 1 was significantly enriched in both the VZ and SVZ/CP of EEDi-treated organoids compared to the DMSO control at protein level (Figure 6I, J).

Taken together, PRC2 inhibition resulted in a loss of H3K27me3 in the three major cell populations of the human developing neocortex, including post-mitotic neurons. Interestingly, the depletion of H3K27me3 did not significantly affect H3K4me2/3 profiles. While a complete loss of the repressive histone mark only resulted in small transcriptomic changes after 5 weeks of EEDi treatment, gene expression analysis suggested that ECM composition may be altered in inhibitor treated organoids. Analysis at protein level for the ECM-related factor Syndecan 1 showed a significant increase in its production upon inhibitor treatment. Our findings, thus, reveal a potential mechanism for non-cell autonomous functions of PRC2 in cortical development, by regulating genes involved in the secretion of signalling molecules and the composition of the ECM in the extracellular niche of the developing neocortex, in which RG, IPs and neurons reside (Figure S7).

## Discussion

Our study employed a highly multiplexed approach to explore a large number of histone modifications at single cell level in the human developing neocortex, providing unprecedented insights into global changes of epigenetic states during human neurogenesis. From this large-scale analysis in human foetal tissue and cortical organoids, PRC2-mediated repressive H3K27me3 emerged as the histone modification that most strongly changed from RG to neurons in the human developing neocortex. Functional experiments revealed that PRC2 regulates the transition of human NPCs from proliferation to differentiation, which is a key determinant of cortical size (Florio and Huttner, 2014). Epigenome analysis identified ECM components and signalling molecules as major targets of PRC2-mediated H3K27me3 in the human developing neocortex, raising the intriguing possibility that PRC2 non-cell autonomously regulates cell fate by modulating the composition of the extracellular stem cell niche in the developing brain.

### Epi-CyTOF represents a large-scale method to detect cell type-specific epigenetic changes

Despite recent advances in single cell epigenome approaches, studying the many layers of histone posttranslational modifications has remained challenging in complex tissues, limiting our understanding of the epigenetic regulation of neural cell fate decisions during neocortex development, especially in human. Typically, global histone modification levels are assessed by Western blotting (Healy et al., 2019; Pasini et al., 2007; Wang et al., 2023), despite the low throughput of the method and limited applicability to tissues that are composed of multiple cell types. CyTOF is a powerful method that takes advantage of metal-conjugated antibodies to quantify proteins within single cells. While well-established in immunology, haematology and oncology (Hartmann and Bendall, 2020; Korin et al., 2018), CyTOF was only recently applied to study chromatin modifications (Cheung et al., 2018; Harpaz et al., 2022). To adopt this powerful method for epigenetic studies of human brain development, we have established a custom Epi-CyTOF antibody panel consisting of >30 epigenetic readouts and 10 neural cell type markers for the identification of major neural cell populations in the human developing neocortex. Following the identification of relevant global changes in histone modifications, epigenomics methods that are applicable for complex tissues, such as cell type-specific CUT&Tag (described here; (Ahanger et al., 2024)) or single cell CUT&Tag (Bartosovic et al., 2021; Meers et al., 2023; Zenk et al., 2024), may be used to gain an in-depth understanding of the epigenetic marking at specific genes.

In the future, Epi-CyTOF can be applied to identify global epigenetic changes in human neurodevelopmental disorders caused by mutations in epigenetic factors (Bolicke and Albert, 2022; Mastrototaro et al., 2017). Moreover, the epigenome is crucial for higher cognitive functions, including memory and learning, and recent evidence implicates the dysregulation of epigenetic mechanisms in neurodegenerative diseases and in aging (Hwang et al., 2017; Wang et al., 2022). Since epigenetic regulation is dynamic and reversible, the epigenome represents a promising target for therapeutic intervention.

### Preservation of epigenetic states in human organoid models of the developing brain

A large number of transcriptome studies have compared gene expression between primary cortical tissue and brain organoids, suggesting that organoids preserve gene regulatory networks and developmental processes, even though metabolic differences were detected (Camp et al., 2015; He et al., 2023; Pollen et al., 2019; Velasco et al., 2019). Less is known about the preservation of epigenomic features in brain organoids. Genome-wide analysis of DNA methylation suggested that most epigenomic features of foetal brain development were recapitulated *in vitro*, even though DNA demethylation occurred in pericentric repeat regions that were not seen in foetal tissue (Luo et al., 2016). Additionally, emerging evidence suggests that chromatin modifications are maintained in forebrain organoids (Amiri et al., 2018; Lewis et al., 2021). Here, we showed that cell type-specific patterns of a large number of histone posttranslational modifications were highly similar in human foetal tissue and human cortical organoids. Moreover, genome-wide distribution of H3K4me2/3 and H3K27me3, two modifications that are centrally involved in cell fate determination (Piunti and Shilatifard, 2016; Schuettengruber et al., 2017), were recapitulated *in vitro*. This represents an important basis for the future application of brain organoid models in the investigation of the epigenetic regulation of human cortical development and neurodevelopmental disorders (Bolicke and Albert, 2022; Mastrototaro et al., 2017).

### PRC2 regulates genes that shape the extracellular environment of the human developing neocortex

Recently, the first studies of Polycomb function in complex organoid models of the human developing brain have revealed a role of PRC2 in regulating the exit from pluripotency (Zenk et al., 2024) and the timing of neuronal maturation (Ciceri et al., 2024), which represents a highly protracted process in humans (Ciceri and Studer, 2024; Wallace and Pollen, 2024). In addition, we show here that PRC2 also regulates the timing during early human cortical development (equivalent to the first trimester of human development), specifically by controlling the neural cell fate progression from RG to IPs to neurons.

Among the most strongly enriched PRC2-target genes, we identified ECM components and signalling molecules, both of which are known to shape the stem cell niche in the human developing neocortex (Massimo and Long, 2021). Based on transcriptomic studies and evolutionary comparisons, we and others have previously proposed that human RG maintain the SVZ as a proliferative niche by locally producing ECM and growth factors to activate self-renewal pathways (Cubillos et al., 2024; Fietz et al., 2012; Florio et al., 2015; Kalebic and Huttner, 2020; Pollen et al., 2015). Here, we found that epigenetic mechanisms, specifically PRC2-mediated H3K27me3, contribute to the regulation of ECM genes. Even though the magnitude of the transcriptional changes was small upon PRC2 inhibition, we did observe changes at protein level for a PRC2-regulated ECM component, Syndecan 1. Such slight changes in the composition of the extracellular environment may thus alter the propensity of NPCs for proliferation, as suggested by evolutionary studies (Cubillos et al., 2024). This provides a mechanistic explanation of previous findings that revealed a role of PRC2 at the global tissue-wide level in the mouse developing brain, rather than a cell-autonomous requirement in neurogenic RG (Amberg et al., 2022).

Overall, our findings suggest that in addition to the cell-autonomous regulation of neurogenic genes, epigenetic regulation of ECM genes may contribute to the determination of neural cell fate in the human developing neocortex, by shaping the composition of the stem cell niche.

## Limitations of study

The Epi-CyTOF approach involves a panel of antibodies directed towards different epigenetic marks. Even though most antibodies against histone posttranslational modifications are well-characterized, cross-reactivities to other modifications have been reported. Independent validation using alternative antibodies or specific inhibitors of epigenetic enzymes (Jin et al., 2022) is recommended for key findings. The cell type marker panel described here is predominantly directed towards different NPC types and neurons. Depending on the scientific question, the panel may be modified to include other cell types that are present in the developing brain, such as endothelial cells, oligodendrocytes and microglia. The suspension mass cytometry technique applied here allows to resolve neural cell types, while spatial information regarding the location of the cells within the tissue is lost. This limitation may be overcome in the future by applying the antibody panel to tissue sections followed by imaging-mass cytometry (IMC)-based analysis (Kuett et al., 2022). Organoids have become an important tool for studies of human brain development, despite known limitations, such as the reduced expansion of the outer SVZ and limited development of neuronal layers. Therefore, we have included primary foetal tissue in this study, which serves as an important reference of the epigenetic state *in vivo*.

## Acknowledgements

We are grateful to the facilities of the CRTD and Dresden Concept partners for the outstanding support provided, notably K. Neumann and her team of the Stem Cell Engineering Facility, H. Hartmann and her team of the Light Microscopy Facility, A. Gompf and her team of the Flow Cytometry Facility, S. Weiche from histology and the wet-lab team at the Dresden Concept Genome Center. We thank all members of the Albert laboratory for help and discussions. We acknowledge the Welcome Trust Sanger Institute (HipSci) for providing the HPSI0114i-kolf_2 iPSC line. M.A. acknowledges funding from the Center for Regenerative Therapies TU Dresden (CRTD), the DFG (Emmy Noether, AL 2231/1-1), the Schram foundation and ERA-NET Neuron (MEPIcephaly; Federal Ministry of Education and Research (BMBF), 01EW2208). The content of this publication is the responsibility of the authors. F.B., A.E., M.O.K. and M.A. acknowledge support from the German Centers for Health Research (DZG). K.S. was supported by the DFG Research Infrastructure NGS_CC (INST 269/468-1 and INST 269/768-1, project 407482635, DcGC DRESDEN-concept Genome Center). This work was supported by the mass cytometry core facility at the CRTD. Ezio Bonifacio (CRTD) raised funding from the DFG (FZT 111) to purchase the suspension mass cytometer CyTOF 2 and from the BMBF to the German Center for Diabetes Research (DZD e.V.) to purchase the suspension mass cytometer CyTOF XT together with the Medical Faculty Carl Gustav Carus, TU Dresden, and the Staatsministerium für Wissenschaft, Kultur und Tourismus (SMWK), Germany.

## Author contributions

Conceptualization, N.D., E.S. and M.A.; Investigation, N.D., E.S., T.M.S., A.N., A.K., S.D. and C.P.; Resources, T.M.S., A.K., K.S., R.P.D, C.B., A.D. and P.W.; Formal Analysis, N.D. and E.S.; Visualization, N.D.; Writing – Original Draft, N.D., E.S. and M.A.; Writing – Review & Editing, N.D., E.S., T.M.S., A.K., C.P. and M.A.; Funding Acquisition, A.E., M.O.K., F.B. and M.A.; Supervision, M.A.

## Declaration of interests

The authors declare no competing interests.

## STAR Methods

### Human foetal tissue

Human foetal brain tissue was obtained from the Department of Gynecology and Obstetrics, University Clinic Carl Gustav Carus of the Technische Universität Dresden, following elective pregnancy termination and informed written maternal consents, and with approval of the local University Hospital Ethical Review Committee (IRB00001473; IORG0001076; ethical approval number EK 355092018), in accordance with the Declaration of Helsinki. The age of foetuses ranged from gestation week (GW) 10 to 14, as assessed by ultrasound measurements of crown-rump length and other standard criteria of developmental stage determination. These developmental time points correspond to an early/mid-neurogenic stage, when the outer SVZ expands, and the production of upper-layer neurons starts. Due to the protection of data privacy, the sex of the human foetuses, from which the human neocortex tissue was obtained, cannot be reported. The sex of the human foetuses is not likely to be of relevance to the results obtained in the present study. The foetal human neocortex tissue samples used in this study reported no health disorders. Foetal human brain tissue was dissected in Tyrode’s solution and used immediately for fixation, processed for CyTOF or snap-frozen for CUT&Tag.

### Human induced pluripotent stem cell lines

All experiments involving hiPSCs were performed in accordance with the ethical standards of the institutional and/or national research committee, as well as with the 1964 Helsinki Declaration, and approved by the University Hospital Ethical Review Committee (IRB00001473; IORG0001076; ethical approval number SR-EK-456092021). Human cortical organoids were generated using the previously generated male human iPSC lines CRTDi004-A (Völkner et al., 2022) and HPSI0114i-kolf_2 (Welcome Trust Sanger Institute, HipSci), derived from healthy donors. The human iPSC lines were maintained on Matrigel-coated (Corning, 354277) culture dishes in mTeSR1 (Stem Cell Technologies, 85850) and passaged using TrypLE Express enzyme (Gibco, 12604021) (CRTDi004-A) or ReLeSR (HPSI0114i- kolf_2) (Stem Cell Technologies, 05873) (Schütze et al., 2022).

### Cortical organoid culture

Human cortical organoids were generated following the sliced neocortical organoid (SNO) protocol, previously described in detail (Cubillos et al., 2024; Qian et al., 2020; Schütze et al., 2022). Briefly, hiPSC colonies (ca. 1.5 mm in diameter) were detached using collagenase (Gibco, 17104019) and transferred to ultra-low attachment six-well plates (Corning, 3471), containing forebrain medium 1, to form embryoid bodies (EBs). EBs were embedded in Matrigel (Corning, 354230) with 20-25 EBs per Matrigel cookie on day 7. The following week (days 7-14), EBs were maintained in forebrain medium 2, with media changes every other day. On day 14, the organoids were released from the Matrigel. Culture is continued in six-well ultra-low attachment plates in forebrain medium 3 on an orbital shaker at 100 rpm. From day 35, forebrain medium 3 is supplemented with 1 % v/v Matrigel. On day 45, organoids were cut on a vibratome into 500-µm-thick slices, to increase supply of oxygen and nutrients throughout the organoids. Therefore, organoids were embedded in 3 % low melting point agarose. Slicing of organoids was repeated at 10 weeks. From day 72, organoids were maintained in forebrain medium 4.

For PRC2 inhibitor treatment, 10 mM EED226 stock solution (MedChemExpress, HY-101117) (Qi et al., 2017) was added to the culture medium to a final concentration of 0.25 µM, 0.5 µM or 5 µM. For the control conditions an equal volume of DMSO (Sigma Aldrich, D2650-100ML) or nothing was added. Medium supplemented with EED226 or DMSO was prepared fresh for each media change. Treatment was started on day 21 of organoid culture and continued until day 56.

### Metal isotope conjugation to antibodies

The antibodies used for the Epi-CyTOF staining were either purchased conjugated with a defined metal isotope or purified within a BSA-free buffer (Table S1). Among those antibodies many were previously validated (Cheung et al., 2018; Harpaz et al., 2022). The conjugation of 100 µg purified antibody with a defined heavy metal tag was done with the MaxPar Antibody Conjugation Kit (Standard BioTools) according to the manufacturer’s recommendations. The protein concentration of the labelled antibodies was measured with a spectrophotometer (Nanodrop 2000c, Thermo Scientific) and adjusted to 0.5 mg/ml with an antibody stabilizer solution (Candor Bioscience, 130050).

### Tissue dissociation for Epi-CyTOF

To prepare the single cell suspensions for the Epi-CyTOF staining, 8-10 cortical organoids (W8) or rice grain sized pieces of human foetal tissue were dissociated using the MACS Neural Tissue Dissociation Kit P (Miltenyi Biotec Inc., 130-092-628) with 6 min of incubation time for enzyme mix 1. Subsequently, the cell pellets were resuspended in 500 μl cold DPBS (Corning, 21-031-CV) and incubated for 10 mins with 1 μl Benzonase (1:500, Sigma, E1014-25KU) to prevent cell clumping and 0.625 μl cisplatin (1:800, Cell-ID™, ^195^Pt, Standard BioTools, 201064) for cell viability staining. From this step onwards, all centrifugation steps were carried out at room temperature (RT). Subsequently, the samples were centrifuged for 7 min at 300 x g and washed with 2 ml Maxpar cell staining buffer (CSB; Standard BioTools, 201068) followed by fixation with 1.6 % paraformaldehyde (PFA, methanol-free, 28906, Thermo Scientific, diluted in DPBS) for 10 min at RT on a rotator. The samples were centrifuged at 900 x g for 10 min and washed twice with 2 ml CSB. Finally, the cell pellets were resuspended in freezing medium (20 % fetal bovine serum (FBS, Sigma, F7524), 10 % dimethyl sulfoxide (DMSO, Sigma, D8418-100ML) in DMEM) and aliquoted (1-2 Mio. cells/ml) before storage store at -70 ^°^C.

### Palladium-based barcoding and cell staining for Epi-CyTOF

For barcoding and staining, the frozen samples were immediately thawed in a water bath at 37 ^°^C and washed with 2 ml CSB. All centrifugation steps are performed at RT. After centrifugation at 900 x g for 10 min, the pellets were resuspended in Maxpar Nuclear Antigen Staining buffer working solution (Mixed 1 part concentrated, 3 parts diluted, Standard BioTools, 201063) and incubated for 30 min at RT for permeabilization. Afterwards, the cells were washed twice by 2 ml Maxpar Nuclear Antigen Staining Perm (NP) buffer (Standard BioTools, 201063), with centrifugations at 900 x g for 10 min. Next, the cell pellets were resuspended in 1 X Fix I Buffer (Standard BioTools, 201065) diluted in PBS and incubated for 10 min at RT, followed by a centrifugation at 900 x g for 10 min. The cells were washed twice with 1 ml 1 X Barcode Perm Buffer (Standard BioTools, 201057). After centrifugation at 900 x g for 10 min, each sample was resuspended in 200 μl 1 X Barcode Perm Buffer. For multiplexing by barcoding, 2.5 μl Palladium barcodes (Cell-ID™ 20-Plex Pd Barcoding Kit, Standard BioTools, 201060) were mixed with 25 μl Barcode Perm buffer, added to each sample separately, and incubated for 30 min at RT. Afterwards, the samples were washed with 2 ml Maxpar CSB and centrifugated at 900 x g for 10 min. The cell pellets were resuspended in 100 μl CSB and pooled in one tube. Up to three million cells were blocked in 50 μl 1 % BSA diluted in NP buffer for 10 min at RT before adding 50 μl pre-mixed and filtered (Merck, UFC30VV00) antibody master mixture in NP. After one hour of staining at RT, the cells were washed with 2 ml CSB, pelleted at 900 x g for 10 min and resuspended in 1 ml Cell-ID™ Intercalator-Iridium (Standard BioTools, 201192A, 1:1000 in Maxpar Fix Perm Buffer). Additional 125 μl 16 % PFA (methanol-free, 28906, Thermo Scientific) were added, and the samples were fixed for 30 min at RT. Finally, the cells were washed with 2 ml CSB, pelleted at 900 x g for 10 min and resuspended in 1 ml freezing medium (20 % FBS and 10 % DMSO in DMEM) to preserve samples at -70 ^°^C until measurement.

### CyTOF measurement

The frozen samples were washed with 2 ml Maxpar cell staining buffer (CSB, Standard BioTools, 201068) and centrifugated at 900 x g for 10 min. The cell pellet was washed with 2 ml Maxpar water (Standard BioTools, 201069), filtered through a 35 µm cell strainer (Falcon, 352235) and counted with the MACSQuant Analyzer (Miltenyi Biotec). For cell acquisition, the cell concentration was adjusted to 6 x 10^5^ cells per ml with Maxpar cell acquisition solution plus (CASplus, Standard BioTools, 201244). As an internal standard for data normalization post-acquisition, 10 % v/v EQ six element calibration beads (Standard BioTools, 201245) with known concentrations of the metal isotopes ^89^Y, ^115^In, ^140^Ce, ^159^Tb, ^175^Lu and ^209^Bi was added. The samples were acquired with the suspension mass cytometer CyTOF2 and CyTOF XT (Standard BioTools) with a constant flow rate of 30 µl and approximately 200 events per second. The recorded linear mode data (LMD) files containing raw ion signals for each push from all available mass channels were processed to FCS file format after event detection, randomization and normalization with the CyTOF software (v9.0.2). The debarcoder function within the software was used to generate separate FCS files per sample based on the 6-choose-3 scheme of each individual Pd code.

### Epi-CyTOF data analysis

After normalization, debarcoding and data cleanup for viable cells, FCS files were uploaded to the cloud-based software OMIQ (Dotmatics) for data analysis. Data were scaled after hyperbolic arcsine (arcsinh) transformation with cofactor 5. For data cleanup, ion clouds, aggregates, and debris were removed through the event length and gaussian parameter. EQ bead signals were removed from the data by cerium isotope ^140^Ce low gating. Cells were identified based on the iridium ^191^Ir and ^193^Ir signal that was also used to remove duplets. Finally, dead cells were excluded by the positive cisplatin ^196^Pt signal to obtain the data of the viable population that was used for downstream analysis. Three main cell type populations were defined according to the expression of SOX2, TBR2, and CTIP2. A standardized gating hierarchy lead to three defined cell clusters. First, all TBR2^+^ events were selected (IPs). Moreover, TBR2^-^ events were further divided into the SOX2^+^/CTIP2^-^ (RG) or CTIP2^+^/SOX2^-^ (neurons) populations. For cluster analysis, the FlowSOM (self.organising maps) algorithm was used to detect consensus metaclusters with a comma-separated k-value of 5 and cluster formation was compared to the hierarchical gating. Uniform Manifold Approximation and Projection (UMAP) was used as the dimensionality reduction algorithm to illustrate the global structure of the 12 samples based on the expression of the 10 cell type-specific markers. The expression level of the epigenetic markers, was illustrated specifically within the clustered data as colour-continuous scatter plots. The median values of each metal tag within the specific cell populations were transformed into log2(FC) and illustrated in heat maps (Graph Pad/Prism software, version 9).

### Immunofluorescence Staining

For immunofluorescence (IF) staining, cortical organoids were removed from culture medium, transferred into 4 % PFA in 120 mM phosphate buffer pH 7.4 and fixed for 30 min at RT. Human foetal tissue was kept in 4 % PFA solution for 24 h at 4 °C. After removing the PFA solution, the tissue was washed twice for 5 min with PBS. Human foetal tissue was subsequently transferred to 15 % Sucrose in PBS, incubated at 4 °C overnight and then transferred to 30 % Sucrose in PBS before storage at 4 °C. Cortical organoids were directly stored in 30 % Sucrose in PBS after fixation and washing. Sucrose-infiltrated tissue was washed in a 1:1 mixture of 30 % Sucrose in PBS and OCT before embedding in the same mixture was performed in cryomolds on dry ice.

Subsequently, 20 µm cryosections were prepared and stored at -20 °C until staining. IF staining was performed as previously described (Florio et al., 2015). Slides containing cryosections were dried for 5 min at RT and briefly washed with PBS before antigen retrieval was performed in 10 mM citrate buffer pH 6.0 for 1 h at 70°C. Following a 5 min PBS wash, the slides were covered with blocking buffer (10 % horse serum and 0.1 % Triton X-100 in PBS) for 30 min at RT. Afterwards the sections were incubated with the primary antibodies diluted in blocking buffer (see dilutions below) overnight at 4°C. The next day, the slides were washed three times for 5 min with PBS, before addition of the secondary antibodies (1:1000) and DAPI (1:1000) diluted in blocking buffer and incubation for 1-2 h at RT. Finally, the slides were washed three times for 5 min with PBS and mounted with Mowiol before long term storage at 4°C.

Images were acquired using a Zeiss ApoTome2 fluorescence microscope with a 20x objective and 1.5- µm thick optical sections. Images are depicted as maximum intensity projections of 10–15 optical sections. Stitching of acquired tile scans was performed using the ZEN software.

### Antibodies dilutions IF

|  |  |
| --- | --- |
| Rat anti-Ctip2 (1:250) | Abcam ab18465 |
| Rat anti-Ki67 (1:500) | Invitrogen 14-5698-82 |
| Goat anti-Sox2 (1:200) | R&D Systems AF2018 |
| Rat anti-PH3 (1:500) | Abcam ab10543 |
| Rabbit anti-TBR2 (1:250) | Abcam ab23345 |
| Rabbit anti-Histone H3K27me3 (1:1600) | Cell Signaling 9733 |
| Rabbit anti-Histone H3K4me2/3 (1:500) | Cell Signaling 9751 |
| Rabbit anti-Ezh2 (1:200) | Cell Signaling 5246 |
| Rabbit anti-Suz12 (1:800) | Cell Signaling 3737 |
| Mouse anti-Zo1 (1:750) | Invitrogen MA3-39100-A647 |
| Rat anti-Tbr2 (1:33) | Invitrogen/EBiosciences 12-4875-82 |
| Rabbit anti-Pax6 (1:300) | BioLegend 901301 |
| Rabbit anti-CC3 (1:400) | Cell Signaling 9661S |

### Image Analysis

All samples were blinded before the acquisition of the data. Quantification was performed using Fiji (Schindelin et al., 2012), processed using Excel (Microsoft), and plotted using Prism (Graphpad Software). Image quantifications were performed either manually or using the Fiji plugin StarDist (Schmidt et al., 2018). Briefly, images were analysed as maximum intensity projections and first cropped to separate the different cortical layers (VZ, SVZ, IZ, CP for foetal human tissue and VZ, SVZ/CP for cortical organoids). Subsequently, all channels were split to use the StarDist 2D plugin in the versatile (fluorescent nuclei) model. If necessary, segmentation was corrected manually. Manual quantifications were performed using the cell counter tool or fluorescence intensity measures in Fiji. For cortical organoids, only ventricles facing the outside of the organoid were analysed, as ventricles in the organoids’ centre might suffer from reduced oxygen and nutrient supply (Qian et al., 2020).

### Isolation of neural cell populations for CUT&Tag and RNA-seq

Whole organoids or rice grain sized pieces of human foetal tissue were transferred to cryovials. After liquids were entirely removed, the vials were snap frozen on dry ice and stored at -70 °C until further processing. The nuclei isolation procedure was modified from a previously described method (Sakib et al., 2021). Using forceps, the frozen tissue was transferred to a microcentrifuge tube and 500 µl Lysis Buffer (10 mM Tris-HCl pH 7.4, 10 mM NaCl, 3 mM MgCl2, 1 % BSA, 0.01 % Tween-20, 0.01 % NP40 Substitute, 0.001 % Digitonin in H2O) were added before homogenising with a pellet pestle (fisher scientific, 12-141-368) 15 times. The homogenised samples were incubated on ice for 2 minutes and mixed 10 times with a wide bore pipet tip followed by 2 more minutes of incubation on ice. With a regular tip, 500 µl chilled wash buffer (10 mM Tris-HCl pH7.4, 10 mM NaCl, 3 mM MgCl2, 1 % BSA, 0.1 % Tween-20 in H2O) were added to each tube and mixed 5 times. The cell suspension was passed through a 70 µm cell strainer (BD Bioscience, 340633) into a LoBind tube followed by additional 300 µl wash buffer. The samples were centrifuged at 500 x g for 5 minutes at 4 °C. The supernatant was removed, and pellets were resuspended in 1 ml of nuclei suspension buffer (NSB; 1 % BSA, 1 X Protease inhibitor EDTA free, 0.1 X RNase inhibitor in PBS). The centrifugation and resuspension steps were repeated once before starting the suspension staining.

For staining, the Alexa Fluor 647 Mouse Anti-Human PAX-6 (BD Pharmingen, 562249, 1:40), PE Mouse Anti-EOMES (BD Pharmingen, 566749, 1:200) and FITC Rat Anti-CTIP2 (abcam, ab123449, 1:500) antibodies were diluted in the nuclei suspension and incubated on a rotator for 1 h at 4 °C in the dark. The stained nuclei suspension was centrifuged for 5 min at 500 x g at 4 °C. Following centrifugation, the supernatant was removed, and the pellet was resuspended in 1 ml NSB. This wash step was repeated once before resuspending in 2 ml NSB, flowing the sample through a 40 µm cell strainer and performing FANS using a BD FACSDiscover S8 Cell Sorter.

IPs were isolated as all TBR2^+^ nuclei. Out of the TBR2^-^ population, CTIP2^+^/PAX6^-^ nuclei were sorted as neurons, while RG were isolated as PAX6^+^/CTIP2^-^. For bulk RNA-seq, 10,000 nuclei were sorted per cell type, while 50,000 nuclei per cell type were used for CUT&Tag.

### Gene expression analysis by RT-qPCR

Gene expression analysis was performed as previously described (Albert et al., 2017; Cubillos et al., 2024). For RT-qPCR, RNA was isolated from 10,000 nuclei (after FANS) using the RNeasy Mini kit (Qiagen, 74104). cDNA was synthesized using random hexamers and Superscript III Reverse Transcriptase (Invitrogen, 18080044) and qPCR was performed with LightCycler 480 SYBR Green I Master Mix (Roche, 4887352001) on a LightCycler 480 (Roche). For each sample, three technical replicates were run. Gene expression data was normalized based on the housekeeping gene *GAPDH*. Primers are listed in Table S4.

### CUT&Tag library preparation

CUT&Tag was performed based on the protocol published previously (Kaya-Okur et al., 2019). The CUT&Tag direct protocol version used was described in detail: dx.doi.org/10.17504/protocols.io.bqwvmxe6. Briefly, 50,000 nuclei per cell type (RG, IP and neurons) were sorted into antibody buffer. Prior to starting the CUT&Tag protocol, the nuclei were centrifuged at 500 x g for 5 min at 4 °C, supernatant was removed and the nuclei were resuspended in 50 µl wash buffer and transferred into PCR tubes. Nuclei were subsequently bound to ConA beads (Polysciences Inc., 86057-3) by adding 3.5 µl ConA beads resuspended in binding buffer to each sample and incubating for 10 min on a rotator at RT.

All following wash steps were performed on a magnetic rack. For blocking of the staining antibodies used for FANS, the supernatant was removed, and the bead-bound nuclei were resuspended in antibody buffer containing 1:1000 Goat F(ab) Anti-Mouse IgG H&L (Abcam, ab6668, RRID: AB_955960) and Goat F(ab) Anti-Rat IgG H&L (Abcam, ab7172). Blocking was performed for 15 min at RT, followed by two wash steps with 100 µl antibody buffer. The bead-bound nuclei were subsequently resuspended in 25 µl antibody buffer containing either 1:50 Rabbit Anti-H3K27me3 (Cell Signaling, 9733) or 1:50 Rabbit Anti-Histone H3K4me2/3 (Cell Signaling, 9751). The primary antibody staining was performed for 1 h on a rotator at RT followed by rotation at 4 °C overnight. After removing the primary antibody, the beads were suspended in 25 µl wash buffer containing 1:100 Guinea Pig Anti-Rabbit IgG (Antibodies-Online, ABIN101961). The secondary antibody was incubated for 45 min on a rotator at RT followed by three washes with 200 µl wash buffer. Pre-loaded pA-Tn5 (Diagenode, C01070001) was diluted 1:250 in 300 buffer and 25 µl were added per sample. After 1 h of incubation on a rotator at RT, samples were washed three times with 200 µl 300-buffer. For tagmentation, the beads were resuspended in 50 µl tagmentation buffer and incubated at 37 °C for 1 h. Finally, the bead-bound nuclei were washed with 50 µl TAPS wash buffer, and resuspended in 5 µl 0.1 % SDS. To release the DNA the samples were incubated at 58 °C for 1 h.

Amplification was performed by addition of 15 µl Triton neutralization solution, 4 µl primer mix (containing 10 µM i5 and i7 uniquely barcoded primers, primers are listed in Table S4) and 25 µl NEBNext High-Fidelity 2X PCR Master Mix (New England Biolabs, M0541S). The following PCR program was run: 5 min 58 °C, 5 min 72 °C, 30 s 98 °C, 13 times [10 s 98 °C, 10 s 60 °C], 1 min 72 °C, hold 8 °C. PCR products were subsequently cleaned up using AMPure XP beads (Beckman Coulter, A63880) at a ratio of 1.3. Sequencing was performed after quantification using a Fragment Analyzer (Agilent) on an Illumina Novaseq 6000 with an average sequencing depth of 20 million fragments.

### CUT&Tag data analysis

CUT&Tag bioinformatics analysis was done using the Nf-core cutandrun pipeline (v3.2.2) (Cheshire et al., 2024; Ewels et al., 2020), with “--genome hg38 --macs-gsize 2805665311 --mito_name ‘chrM’ -- normalization_binsize 1 --use_control ‘false’ --dedup_target_reads --peakcaller SEACR -- seacr_peak_threshold 0.01” and default settings otherwise. The blacklist provided was derived from (Amemiya et al., 2019). The pipeline automatically runs all required steps on a library-level, including (but not limited to) raw read quality control, adapter trimming, alignment, filtering, generation of coverage tracks, peak calling and peak quality control. Finally, a consensus peak set is created across all libraries (--consensus_peak_mode all) or libraries from the same cell type (--consensus_peak_mode group). H3K27me3 and H3K4me2/3 CUT&Tag data was processed separately. All datasets acquired from human foetal tissue were CPM normalized (--normalisation_mode CPM). Data acquired from inhibitor and DMSO treated cortical organoids was used unnormalized (--normalisation_mode None), as CPM normalization was not appropriate for portraying the strong differences expected between the two analysed conditions. For direct comparison of the two histone marks, consensus peak sets generated within the nf-core pipeline were used.

The nf-core pipeline output for each histone modification was further processed with the R package Diffbind (v 3.12.0) (Stark and Brown, 2011) for visualization and differential analysis. Raw reads and called peaks were imported and counts were generated within previously determined consensus peak regions without centering the summits. Generated counts for human foetal tissue data were normalized by library size while counts for the cortical organoid data were used unnormalized due to the expected differences in library size. Regions of differential enrichment were determined with DESeq2 (v1.42.1) (Love et al., 2014) (p-adjust < 0.05; logFC > 1). Heatmaps and profile plots displaying histone mark enrichment were generated using deeptools (v3.5.1) (Ramirez et al., 2016). Regions identified to be enriched for a histone modification were annotated to genomic features and the nearest neighbouring gene using the R packages ChIPseeker (v1.38.0) (Yu et al., 2015) and TxDb.Hsapiens.UCSC.hg38.knownGene (v3.18.0) (Team and Maintainer, 2019). Pathway analysis for those identified genes was performed using ReactomePA (v1.46.0) analysis (Yu and He, 2016).

### RNA-seq library preparation

Transcriptome libraries were prepared using the SmartSeq-2 protocol (Picelli et al., 2013), as previously described (Cubillos et al., 2024). Isolated total RNA from an equivalent of 10,000 nuclei was denatured for 3 min at 72 °C in 4 μl hypotonic buffer (0.2 % Triton-X 100) in the presence of 2.5 mM dNTP, 100 nM dT-primer and 4 U RNase Inhibitor (Promega, N2611). Reverse transcription was performed at 42 °C for 90 min after filling up to 10 µl with RT buffer mix for a final concentration of 1X Superscript II buffer (Invitrogen, 18064022), 1 M betaine, 5 mM DTT, 6 mM MgCl2, 1 µM TSO-primer (Table S4), 9 U RNase inhibitor, and 90 U Superscript II. The reverse transcriptase was inactivated at 70 °C for 15 min. For subsequent PCR amplification of the cDNA the optimal PCR cycle number was determined with an aliquot of 1 μl unpurified cDNA in a 10 μl qPCR containing 1X KAPA HiFi Hotstart Readymix (Roche, 07958927001), 1X SYBR Green, and 0.2 μM UP-primer (Table S4). The residual 9 μl cDNA were subsequently amplified using KAPA HiFi HotStart Readymix (Roche, 07958927001) at a 1X concentration together with 250 nM UP-primer under the following cycling conditions: initial denaturation at 98 °C for 3 min, 12 cycles [98 °C 20 s, 67 °C 15 s, 72 °C 6 min] and final elongation at 72 °C for 5 min. Amplified cDNA was purified using 1X volume of Sera-Mag SpeedBeads (GE Healthcare) resuspended in a buffer consisting of 10 mM Tris, 20 mM EDTA, 18.5% (w/v) PEG 8000, and 2 M sodium chloride solution. The cDNA quality and concentration were determined using a Fragment Analyzer (Agilent).

For library preparation, 2 µl amplified cDNA was tagmented in 1X Tagmentation Buffer using 0.8 µl bead-linked transposome (Illumina DNA Prep, (M) Tagmentation, Illumina) at 55 °C for 15 min in a total volume of 4 µl. The reaction was stopped by adding 1 µl of 0.1 % SDS (37 °C, 15 min). Magnetic beads were bound to a magnet, the supernatant was removed, beads were resuspended in 14 µl indexing PCR Mix containing 1X KAPA Hifi HotStart ReadyMix (Roche) and 700 nM unique dual indexing primers (i5 and i7), and subjected to a PCR (72 °C 3 min, 98 °C 30 s, 12 cycles [98 °C 10 s, 63 °C 20 s, 72 °C 1 min], and 72 °C 5 min). Libraries were purified with 0.9X volume Sera-Mag SpeedBeads, followed by a double size selection with 0.6 X and 0.9 X volume of beads. Sequencing was performed after quantification using a Fragment Analyzer on an Illumina Novaseq 6000 with an average sequencing depth of 60 million fragments.

### RNA-seq data analysis

RNA-seq data analysis was performed as previously described (Cubillos et al., 2024; Hoffmann et al., 2024). Quality control of the sequencing data was performed with FastQC (v0.11.9). Kallisto (v0.64.1) (Bray et al., 2016) was used to align trimmed reads to mouse GRChg38. For further processing, data was imported into R using Tximport (v1.30.0) (Soneson et al., 2015) and EnsDb.Hsapiens.v86 (v2.99.0) (Rainer, 2017). Raw fragment normalization based on library size and testing for differential expression between genotypes was performed with edgeR package (v4.0.16) (Robinson et al., 2010) with a false discovery rate (FDR) of 5 % and a Log2 fold change threshold of 0.5. Volcano plots and heatmaps were generated with the R-packages Ggplot2 (v3.1.3.1) (Wickham et al., 2016). Differentially expressed genes were analysed by gene ontology (https://geneontology.org/) (Ashburner et al., 2000; Thomas et al., 2022) analysis.

### Statistical analysis

Sample sizes are reported in each figure legend. Sample sizes were estimated based on previous literature (Albert et al., 2017; Cubillos et al., 2024; Schmitz et al., 2011). All statistical analysis was performed using Prism (GraphPad Software). Normal distribution of datasets was tested by Shapiro-Wilk or Kolmogorov-Smirnov test. The tests used are indicated in the figure legend for each quantification. Significant changes are indicated by stars for each graph and described in the figure legends.

## Notes

### Competing Interest Statement

The authors have declared no competing interest.

## References

Ahanger, S.H., Zhang, C., Semenza, E.R., Gil, E., Cole, M.A., Wang, L., Kriegstein, A.R., and Lim, D.A. (2024). Spatial 3D genome organization controls the activity of bivalent chromatin during human neurogenesis. bioRxiv.

Albert, M., and Huttner, W.B. (2018). Epigenetic and Transcriptional Pre-patterning-An Emerging Theme in Cortical Neurogenesis. Front Neurosci 12, 359.

Albert, M., Kalebic, N., Florio, M., Lakshmanaperumal, N., Haffner, C., Brandl, H., Henry, I., and Huttner, W.B. (2017). Epigenome profiling and editing of neocortical progenitor cells during development. EMBO J 36, 2642–2658.

Amberg, N., Laukoter, S., and Hippenmeyer, S. (2019). Epigenetic cues modulating the generation of cell-type diversity in the cerebral cortex. J Neurochem 149, 12–26.

Amberg, N., Pauler, F.M., Streicher, C., and Hippenmeyer, S. (2022). Tissue-wide genetic and cellular landscape shapes the execution of sequential PRC2 functions in neural stem cell lineage progression. Sci Adv 8, eabq1263.

Amemiya, H.M., Kundaje, A., and Boyle, A.P. (2019). The ENCODE Blacklist: Identification of Problematic Regions of the Genome. Sci Rep 9, 9354.

Amiri, A., Coppola, G., Scuderi, S., Wu, F., Roychowdhury, T., Liu, F., Pochareddy, S., Shin, Y., Safi, A., Song, L., et al. (2018). Transcriptome and epigenome landscape of human cortical development modeled in organoids. Science 362.

Ashburner, M., Ball, C.A., Blake, J.A., Botstein, D., Butler, H., Cherry, J.M., Davis, A.P., Dolinski, K., Dwight, S.S., Eppig, J.T., et al. (2000). Gene ontology: tool for the unification of biology. The Gene Ontology Consortium. Nat Genet 25, 25–29.

Barth, T.K., and Imhof, A. (2010). Fast signals and slow marks: the dynamics of histone modifications. Trends Biochem Sci 35, 618–626.

Bartosovic, M., Kabbe, M., and Castelo-Branco, G. (2021). Single-cell CUT&Tag profiles histone modifications and transcription factors in complex tissues. Nat Biotechnol 39, 825–835.

Bolicke, N., and Albert, M. (2022). Polycomb-mediated gene regulation in human brain development and neurodevelopmental disorders. Dev Neurobiol 82, 345–363.

Borrell, V., and Reillo, I. (2012). Emerging roles of neural stem cells in cerebral cortex development and evolution. Dev Neurobiol 72, 955–971.

Boyer, L.A., Plath, K., Zeitlinger, J., Brambrink, T., Medeiros, L.A., Lee, T.I., Levine, S.S., Wernig, M., Tajonar, A., Ray, M.K., et al. (2006). Polycomb complexes repress developmental regulators in murine embryonic stem cells. Nature 441, 349–353.

Bray, N.L., Pimentel, H., Melsted, P., and Pachter, L. (2016). Near-optimal probabilistic RNA-seq quantification. Nat Biotechnol 34, 525–527.

Camp, J.G., Badsha, F., Florio, M., Kanton, S., Gerber, T., Wilsch-Brauninger, M., Lewitus, E., Sykes, A., Hevers, W., Lancaster, M., et al. (2015). Human cerebral organoids recapitulate gene expression programs of fetal neocortex development. Proc Natl Acad Sci U S A 112, 15672–15677.

Cheshire, C., Charlotte-west, Rönkkö, T., bot, n.-c., Patel, H., Tamara-hodgetts, Ladd, D., Thiery, A., Fields, C., Deu-Pons, J., et al. (2024). nf-core/cutandrun v3.2.2 Iridium Ibis.

Cheung, P., Vallania, F., Warsinske, H.C., Donato, M., Schaffert, S., Chang, S.E., Dvorak, M., Dekker, C.L., Davis, M.M., Utz, P.J., et al. (2018). Single-Cell chromatin modification profiling reveals increased epigenetic variations with aging. Cell 173, 1385–1397 e1314.

Ciceri, G., Baggiolini, A., Cho, H.S., Kshirsagar, M., Benito-Kwiecinski, S., Walsh, R.M., Aromolaran, K.A., Gonzalez-Hernandez, A.J., Munguba, H., Koo, S.Y., et al. (2024). An epigenetic barrier sets the timing of human neuronal maturation. Nature 626, 881–890.

Ciceri, G., and Studer, L. (2024). Epigenetic control and manipulation of neuronal maturation timing. Curr Opin Genet Dev 85, 102164.

Corley, M., and Kroll, K.L. (2015). The roles and regulation of Polycomb complexes in neural development. Cell Tissue Res 359, 65–85.

Cubillos, P., Ditzer, N., Kolodziejczyk, A., Schwenk, G., Hoffmann, J., Schutze, T.M., Derihaci, R.P., Birdir, C., Kollner, J.E., Petzold, A., et al. (2024). The growth factor EPIREGULIN promotes basal progenitor cell proliferation in the developing neocortex. EMBO J 43, 1388–1419.

den Hoed, J., Sollis, E., Venselaar, H., Estruch, S.B., Deriziotis, P., and Fisher, S.E. (2018). Functional characterization of TBR1 variants in neurodevelopmental disorder. Sci Rep 8, 14279.

Deriziotis, P., O’Roak, B.J., Graham, S.A., Estruch, S.B., Dimitropoulou, D., Bernier, R.A., Gerdts, J., Shendure, J., Eichler, E.E., and Fisher, S.E. (2014). De novo TBR1 mutations in sporadic autism disrupt protein functions. Nat Commun 5, 4954.

Ewels, P.A., Peltzer, A., Fillinger, S., Patel, H., Alneberg, J., Wilm, A., Garcia, M.U., Di Tommaso, P., and Nahnsen, S. (2020). The nf-core framework for community-curated bioinformatics pipelines. Nat Biotechnol 38, 276–278.

Fietz, S.A., Lachmann, R., Brandl, H., Kircher, M., Samusik, N., Schroder, R., Lakshmanaperumal, N., Henry, I., Vogt, J., Riehn, A., et al. (2012). Transcriptomes of germinal zones of human and mouse fetal neocortex suggest a role of extracellular matrix in progenitor self-renewal. Proc Natl Acad Sci U S A 109, 11836–11841.

Florio, M., Albert, M., Taverna, E., Namba, T., Brandl, H., Lewitus, E., Haffner, C., Sykes, A., Wong, F.K., Peters, J., et al. (2015). Human-specific gene ARHGAP11B promotes basal progenitor amplification and neocortex expansion. Science 347, 1465–1470.

Florio, M., and Huttner, W.B. (2014). Neural progenitors, neurogenesis and the evolution of the neocortex. Development 141, 2182–2194.

Giandomenico, S.L., and Lancaster, M.A. (2017). Probing human brain evolution and development in organoids. Curr Opin Cell Biol 44, 36–43.

Harpaz, N., Mittelman, T., Beresh, O., Griess, O., Furth, N., Salame, T.M., Oren, R., Fellus-Alyagor, L., Harmelin, A., Alexandrescu, S., et al. (2022). Single-cell epigenetic analysis reveals principles of chromatin states in H3.3-K27M gliomas. Mol Cell 82, 2696–2713 e2699.

Hartmann, F.J., and Bendall, S.C. (2020). Immune monitoring using mass cytometry and related high-dimensional imaging approaches. Nat Rev Rheumatol 16, 87–99.

He, Z., Dony, L., Fleck, J.S., Szałata, A., Li, K.X., Slišković, I., Lin, H., Santel, M., Atamian, A., Quadrato, G., et al. (2023). An integrated transcriptomic cell atlas of human neural organoids. bioRxiv, 2023.2010.2005.561097.

Healy, E., Mucha, M., Glancy, E., Fitzpatrick, D.J., Conway, E., Neikes, H.K., Monger, C., Van Mierlo, G., Baltissen, M.P., Koseki, Y., et al. (2019). PRC2.1 and PRC2.2 Synergize to Coordinate H3K27 Trimethylation. Mol Cell 76, 437–452 e436.

Hirabayashi, Y., and Gotoh, Y. (2010). Epigenetic control of neural precursor cell fate during development. Nat Rev Neurosci 11, 377–388.

Hoffmann, J., Schütze, T.M., Kolodziejczyk, A., Kränkel, A., Reinhardt, S., Derihaci, R.P., Birdir, C., Wimberger, P., Koseki, H., and Albert, M. (2024). Canonical and non-canonical PRC1 differentially contribute to the regulation of neural stem cell fate. bioRxiv, 2024.2008.2007.606990.

Hoye, M.L., and Silver, D.L. (2020). Decoding mixed messages in the developing cortex: translational regulation of neural progenitor fate. Curr Opin Neurobiol 66, 93–102.

Hwang, J.Y., Aromolaran, K.A., and Zukin, R.S. (2017). The emerging field of epigenetics in neurodegeneration and neuroprotection. Nat Rev Neurosci 18, 347–361.

Jin, Y., Liu, T., Luo, H., Liu, Y., and Liu, D. (2022). Targeting Epigenetic Regulatory Enzymes for Cancer Therapeutics: Novel Small-Molecule Epidrug Development. Front Oncol 12, 848221.

Kalebic, N., and Huttner, W.B. (2020). Basal progenitor morphology and neocortex evolution. Trends Neurosci 43, 843–853.

Kaya-Okur, H.S., Wu, S.J., Codomo, C.A., Pledger, E.S., Bryson, T.D., Henikoff, J.G., Ahmad, K., and Henikoff, S. (2019). CUT&Tag for efficient epigenomic profiling of small samples and single cells. Nat Commun 10, 1930.

Korin, B., Dubovik, T., and Rolls, A. (2018). Mass cytometry analysis of immune cells in the brain. Nat Protoc 13, 377–391.

Kuett, L., Catena, R., Ozcan, A., Pluss, A., Cancer Grand Challenges, I.C., Schraml, P., Moch, H., de Souza, N., and Bodenmiller, B. (2022). Three-dimensional imaging mass cytometry for highly multiplexed molecular and cellular mapping of tissues and the tumor microenvironment. Nat Cancer 3, 122–133.

Lewis, E.M.A., Kaushik, K., Sandoval, L.A., Antony, I., Dietmann, S., and Kroll, K.L. (2021). Epigenetic regulation during human cortical development: Seq-ing answers from the brain to the organoid. Neurochem Int 147, 105039.

Love, M.I., Huber, W., and Anders, S. (2014). Moderated estimation of fold change and dispersion for RNA-seq data with DESeq2. Genome Biol 15, 550.

Lui, J.H., Hansen, D.V., and Kriegstein, A.R. (2011). Development and evolution of the human neocortex. Cell 146, 18–36.

Luo, C., Lancaster, M.A., Castanon, R., Nery, J.R., Knoblich, J.A., and Ecker, J.R. (2016). Cerebral organoids recapitulate epigenomic signatures of the human fetal brain. Cell Rep 17, 3369–3384.

Massimo, M., and Long, K.R. (2021). Orchestrating human neocortex development across the scales; from micro to macro. Semin Cell Dev Biol.

Mastrototaro, G., Zaghi, M., and Sessa, A. (2017). Epigenetic Mistakes in Neurodevelopmental Disorders. J Mol Neurosci 61, 590–602.

Meers, M.P., Llagas, G., Janssens, D.H., Codomo, C.A., and Henikoff, S. (2023). Multifactorial profiling of epigenetic landscapes at single-cell resolution using MulTI-Tag. Nat Biotechnol 41, 708–716.

Millan-Zambrano, G., Burton, A., Bannister, A.J., and Schneider, R. (2022). Histone post-translational modifications - cause and consequence of genome function. Nat Rev Genet 23, 563–580.

Moore, S.W., Tessier-Lavigne, M., and Kennedy, T.E. (2007). Netrins and their receptors. Adv Exp Med Biol 621, 17–31.

Mouthon, M.A., Morizur, L., Dutour, L., Pineau, D., Kortulewski, T., and Boussin, F.D. (2020). Syndecan-1 Stimulates Adult Neurogenesis in the Mouse Ventricular-Subventricular Zone after Injury. iScience 23, 101784.

Nano, P.R., and Bhaduri, A. (2022). Evaluation of advances in cortical development using model systems. Dev Neurobiol 82, 408–427.

Noack, F., Vangelisti, S., Ditzer, N., Chong, F., Albert, M., and Bonev, B. (2023). Joint epigenome profiling reveals cell-type-specific gene regulatory programmes in human cortical organoids. Nat Cell Biol 25, 1873–1883.

O’Carroll, D., Erhardt, S., Pagani, M., Barton, S.C., Surani, M.A., and Jenuwein, T. (2001). The polycomb-group gene Ezh2 is required for early mouse development. Mol Cell Biol 21, 4330–4336.

Pasca, S.P. (2018). The rise of three-dimensional human brain cultures. Nature 553, 437–445.

Pasini, D., Bracken, A.P., Hansen, J.B., Capillo, M., and Helin, K. (2007). The polycomb group protein Suz12 is required for embryonic stem cell differentiation. Mol Cell Biol 27, 3769–3779.

Pasini, D., Bracken, A.P., Jensen, M.R., Lazzerini Denchi, E., and Helin, K. (2004). Suz12 is essential for mouse development and for EZH2 histone methyltransferase activity. EMBO J 23, 4061–4071.

Pereira, J.D., Sansom, S.N., Smith, J., Dobenecker, M.W., Tarakhovsky, A., and Livesey, F.J. (2010). Ezh2, the histone methyltransferase of PRC2, regulates the balance between self-renewal and differentiation in the cerebral cortex. Proc Natl Acad Sci U S A 107, 15957–15962.

Picelli, S., Bjorklund, A.K., Faridani, O.R., Sagasser, S., Winberg, G., and Sandberg, R. (2013). Smart- seq2 for sensitive full-length transcriptome profiling in single cells. Nat Methods 10, 1096–1098.

Piunti, A., and Shilatifard, A. (2016). Epigenetic balance of gene expression by Polycomb and COMPASS families. Science 352, aad9780.

Pollen, A.A., Bhaduri, A., Andrews, M.G., Nowakowski, T.J., Meyerson, O.S., Mostajo-Radji, M.A., Di Lullo, E., Alvarado, B., Bedolli, M., Dougherty, M.L., et al. (2019). Establishing Cerebral Organoids as Models of Human-Specific Brain Evolution. Cell 176, 743–756 e717.

Pollen, A.A., Kilik, U., Lowe, C.B., and Camp, J.G. (2023). Human-specific genetics: new tools to explore the molecular and cellular basis of human evolution. Nat Rev Genet, 1-25.

Pollen, A.A., Nowakowski, T.J., Chen, J., Retallack, H., Sandoval-Espinosa, C., Nicholas, C.R., Shuga, J., Liu, S.J., Oldham, M.C., Diaz, A., et al. (2015). Molecular identity of human outer radial glia during cortical development. Cell 163, 55–67.

Qi, W., Zhao, K., Gu, J., Huang, Y., Wang, Y., Zhang, H., Zhang, M., Zhang, J., Yu, Z., Li, L., et al. (2017). An allosteric PRC2 inhibitor targeting the H3K27me3 binding pocket of EED. Nat Chem Biol 13, 381–388.

Qian, X., Su, Y., Adam, C.D., Deutschmann, A.U., Pather, S.R., Goldberg, E.M., Su, K., Li, S., Lu, L., Jacob, F., et al. (2020). Sliced Human Cortical Organoids for Modeling Distinct Cortical Layer Formation. Cell Stem Cell 26, 766–781 e769.

Rainer, J. (2017). EnsDb.Hsapiens.v86: Ensembl based annotation package.

Ramirez, F., Ryan, D.P., Gruning, B., Bhardwaj, V., Kilpert, F., Richter, A.S., Heyne, S., Dundar, F., and Manke, T. (2016). deepTools2: a next generation web server for deep-sequencing data analysis. Nucleic Acids Res 44, W160–165.

Robinson, M.D., McCarthy, D.J., and Smyth, G.K. (2010). edgeR: a Bioconductor package for differential expression analysis of digital gene expression data. Bioinformatics 26, 139–140.

Sakib, M.S., Sokpor, G., Nguyen, H.P., Fischer, A., and Tuoc, T. (2021). Intranuclear immunostaining-based FACS protocol from embryonic cortical tissue. STAR Protoc 2, 100318.

Schindelin, J., Arganda-Carreras, I., Frise, E., Kaynig, V., Longair, M., Pietzsch, T., Preibisch, S., Rueden, C., Saalfeld, S., Schmid, B., et al. (2012). Fiji: an open-source platform for biological-image analysis. Nat Methods 9, 676–682.

Schmidt, U., Weigert, M., Broaddus, C., and Myers, G. (2018). Cell Detection with Star-Convex Polygons (Cham: Springer International Publishing).

Schmitz, S.U., Albert, M., Malatesta, M., Morey, L., Johansen, J.V., Bak, M., Tommerup, N., Abarrategui, I., and Helin, K. (2011). Jarid1b targets genes regulating development and is involved in neural differentiation. EMBO J 30, 4586–4600.

Schuettengruber, B., Bourbon, H.M., Di Croce, L., and Cavalli, G. (2017). Genome Regulation by Polycomb and Trithorax: 70 Years and Counting. Cell 171, 34–57.

Schütze, T.M., Bölicke, N., Sameith, K., and Albert, M. (2022). Profiling Cell Type-Specific Gene Regulatory Regions in Human Cortical Organoids. In Brain Organoid Research, J. Gopalakrishnan, ed. (New York, NY: Springer US), pp. 17–41.

Soneson, C., Love, M.I., and Robinson, M.D. (2015). Differential analyses for RNA-seq: transcript-level estimates improve gene-level inferences. F1000Res 4, 1521.

Stark, R., and Brown, G. (2011). DiffBind: differential binding analysis of ChIP-Seq peak data.

Team, B.C., and Maintainer, B.P. (2019). TxDb.Hsapiens.UCSC.hg38.knownGene: Annotation package for TxDb object(s).

Telley, L., Agirman, G., Prados, J., Amberg, N., Fievre, S., Oberst, P., Bartolini, G., Vitali, I., Cadilhac, C., Hippenmeyer, S., et al. (2019). Temporal patterning of apical progenitors and their daughter neurons in the developing neocortex. Science 364.

Thomas, P.D., Ebert, D., Muruganujan, A., Mushayahama, T., Albou, L.P., and Mi, H. (2022). PANTHER: Making genome-scale phylogenetics accessible to all. Protein Sci 31, 8–22.

Tsuboi, M., Hirabayashi, Y., and Gotoh, Y. (2019). Diverse gene regulatory mechanisms mediated by Polycomb group proteins during neural development. Curr Opin Neurobiol 59, 164–173.

Turner, B.M. (1993). Decoding the nucleosome. Cell 75, 5–8.

Vanderhaeghen, P., and Polleux, F. (2023). Developmental mechanisms underlying the evolution of human cortical circuits. Nat Rev Neurosci 24, 213–232.

Velasco, S., Kedaigle, A.J., Simmons, S.K., Nash, A., Rocha, M., Quadrato, G., Paulsen, B., Nguyen, L., Adiconis, X., Regev, A., et al. (2019). Individual brain organoids reproducibly form cell diversity of the human cerebral cortex. Nature 570, 523–527.

Völkner, M., Wagner, F., Steinheuer, L.M., Carido, M., Kurth, T., Yazbeck, A., Schor, J., Wieneke, S., Ebner, L.J.A., Del Toro Runzer, C., et al. (2022). HBEGF-TNF induce a complex outer retinal pathology with photoreceptor cell extrusion in human organoids. Nat Commun 13, 6183.

Wallace, J.L., and Pollen, A.A. (2024). Human neuronal maturation comes of age: cellular mechanisms and species differences. Nat Rev Neurosci 25, 7–29.

Wang, H., Fan, Z., Shliaha, P.V., Miele, M., Hendrickson, R.C., Jiang, X., and Helin, K. (2023). H3K4me3 regulates RNA polymerase II promoter-proximal pause-release. Nature 615, 339–348.

Wang, K., Liu, H., Hu, Q., Wang, L., Liu, J., Zheng, Z., Zhang, W., Ren, J., Zhu, F., and Liu, G.H. (2022). Epigenetic regulation of aging: implications for interventions of aging and diseases. Signal Transduct Target Ther 7, 374.

Wang, Q., Yang, L., Alexander, C., and Temple, S. (2012). The niche factor syndecan-1 regulates the maintenance and proliferation of neural progenitor cells during mammalian cortical development. PLoS One 7, e42883.

Wickham, H., Navarro, D., and Pedersen, T.L. (2016). ggplot2: Elegant Graphics for Data Analysis (3e), Third edition edn (Springer).

Yao, B., Christian, K.M., He, C., Jin, P., Ming, G.L., and Song, H. (2016). Epigenetic mechanisms in neurogenesis. Nat Rev Neurosci 17, 537–549.

Yu, G., and He, Q.Y. (2016). ReactomePA: an R/Bioconductor package for reactome pathway analysis and visualization. Mol Biosyst 12, 477–479.

Yu, G., Wang, L.G., and He, Q.Y. (2015). ChIPseeker: an R/Bioconductor package for ChIP peak annotation, comparison and visualization. Bioinformatics 31, 2382–2383.

Zenk, F., Fleck, J.S., Jansen, S.M.J., Kashanian, B., Eisinger, B., Santel, M., Dupre, J.S., Camp, J.G., and Treutlein, B. (2024). Single-cell epigenomic reconstruction of developmental trajectories from pluripotency in human neural organoid systems. Nat Neurosci 27, 1376–1386.

