## Supplementary Figures for "Epigenome profiling identifies H3K27me3 regulation of extra-cellular matrix composition in human corticogenesis"

### SUPPLEMENTARY INFORMATION

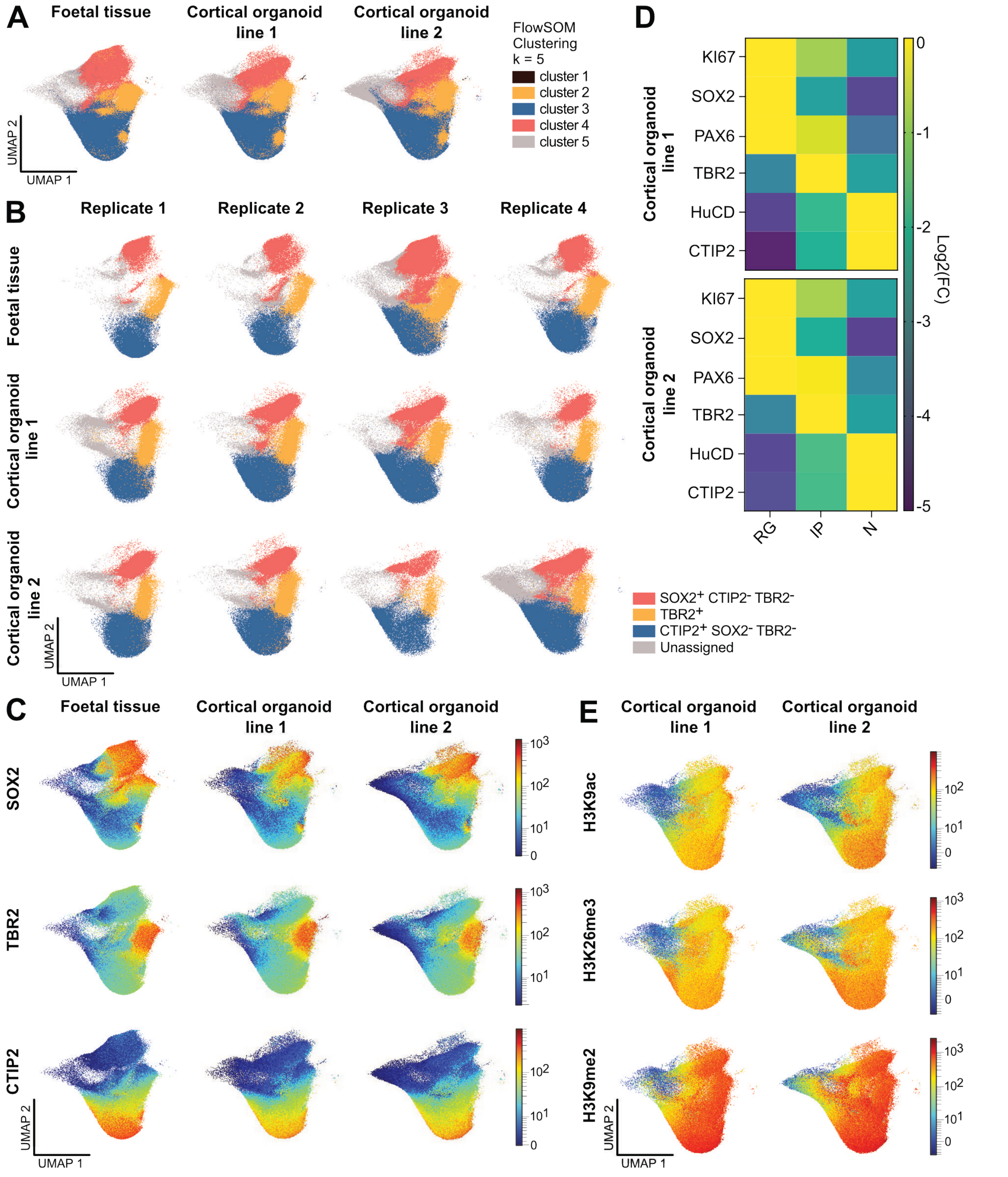


**Figure S1 related to Figure 1. Epi-CyTOF single cell analysis of the human developing neocortex.**

(A) Unbiased identification of cortical cell populations by FlowSOM clustering (k=5) using cell type markers results in similar clusters as identified by gating strategy based on SOX2, TBR2 and CTIP2. Clustering is shown for foetal tissue and cortical organoids derived from two independent iPSC lines. (B) UMAP analysis of cell type clustering for 4 replicates of human foetal tissue (GW12–14; from two tissue samples) and 4 replicates of cortical organoids each from two independent iPSC lines (8W; 4 batches each). (C) UMAP colour-continuous scatter plots for the RG marker SOX2, the IP marker TBR2 and the neuronal marker CTIP2 for human foetal tissue and cortical organoids derived from two independent iPSC lines (combination of 4 replicates each). Scale represents metal isotope tag intensity. (D) Heat map of Log2 fold changes of median metal tag intensities for cell type markers in RG, IP and N from cortical organoids derived from two independent iPSC lines. (E) UMAP colour-continuous scatter plots for the intensity of H3K9ac, H3K36me3 and H3K9me2 in cortical organoids from iPSC line 1 and 2.

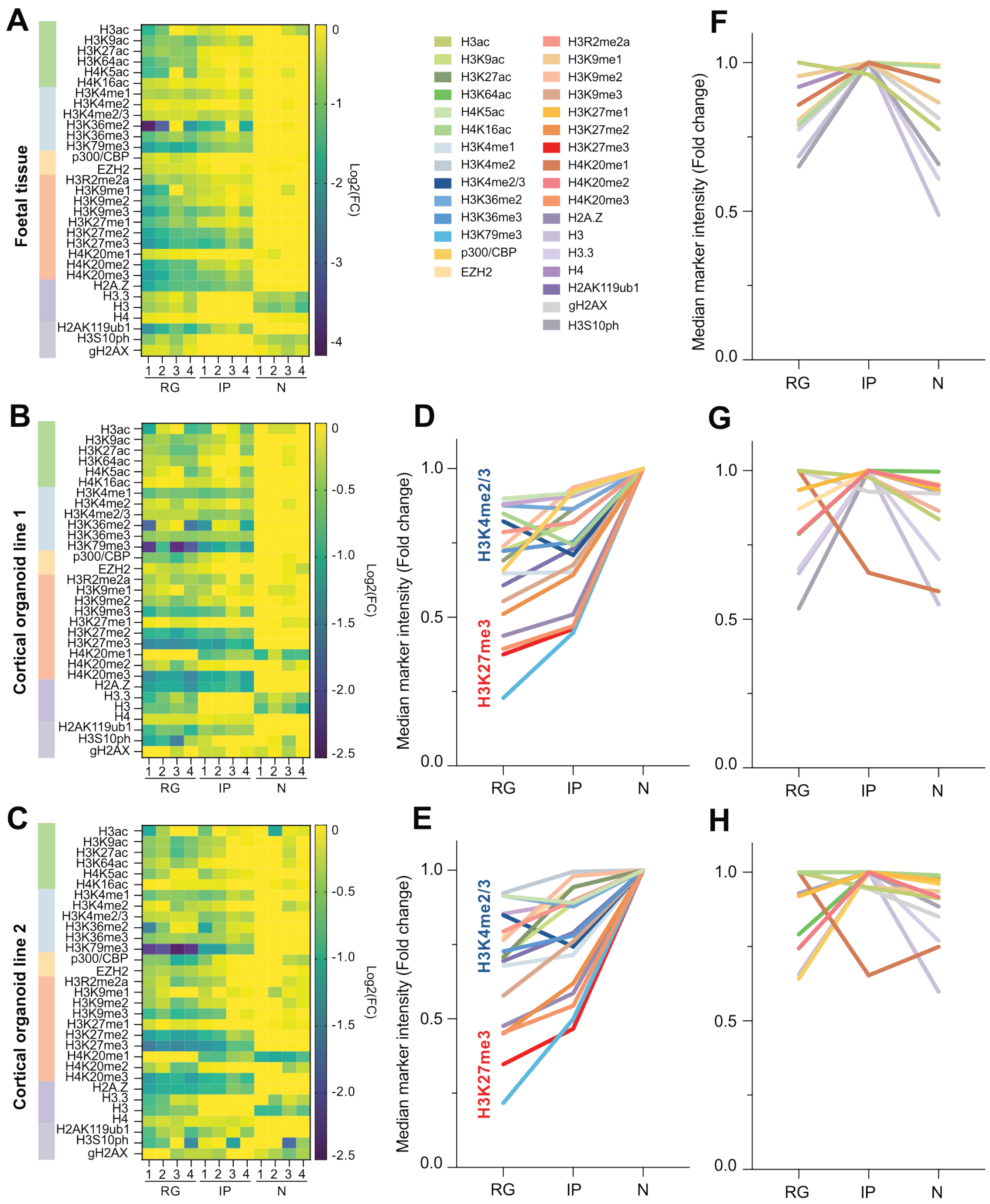

**Figure S2 related to Figure 2. Global levels of 31 epigenetic marks in the human developing neocortex detected by Epi-CyTOF.**

(A–C) Heat maps of Log2 fold changes of median metal tag intensities for epigenetics markers in RG, IP and N for human foetal tissue (GW12-14) (A) and cortical organoids (W8) from iPSC line 1 (B) and line 2 (C). All 4 replicates are displayed individually. The order of the marks is according to the Epi-CyTOF antibody panel with active acetylation (green), active methylation (blue), enzymes (orange), repressive methylation (red), histone variants (purple) and other marks (grey). (D, E) Line plots displaying epigenetic marks with increasing abundance from RG to N. Values represent fold change of Epi-CyTOF median metal tag intensities of cortical organoids from iPSC line 1 (D) and line 2 (E) (combination of 4 replicates each). (F–H) Line plots displaying remaining epigenetic marks for human foetal tissue (F) and cortical organoids from iPSC line 1 (G) and line 2 (H).

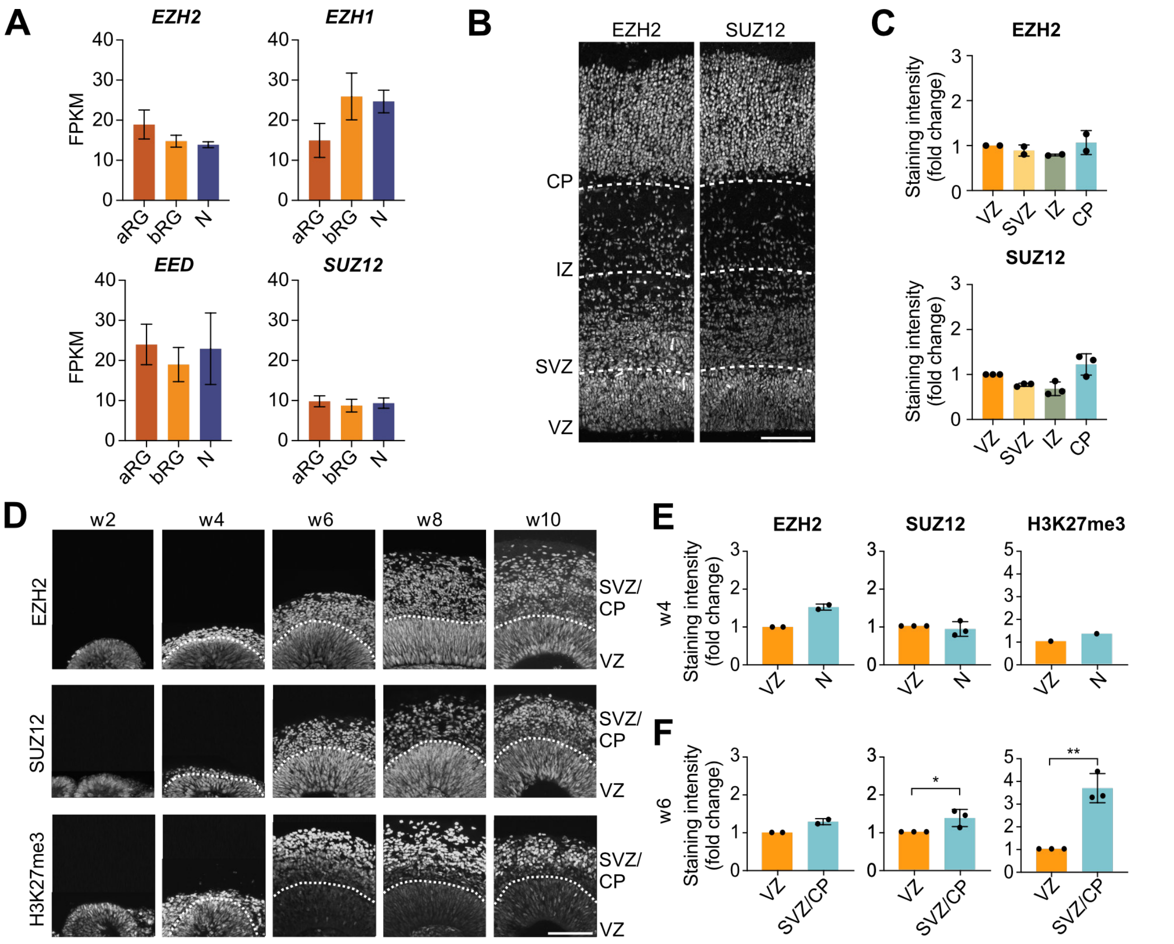


**Figure S3 related to Figure 2. Expression of PRC2 and H3K27me3 marking in the human developing neocortex**.
(A) mRNA levels of PRC2 components in aRG, bRG and N from human foetal cortex analysed by RNA-seq (data from (Florio et al., 2015)). (B) Immunofluorescence staining for EZH2 and SUZ12 of human foetal tissue (GW12–14). (C) Quantification of EZH2 and SUZ12 intensity in nuclei (segmented based on DAPI) in the VZ, SVZ, IZ and CP relative to the VZ. (D) Immunofluorescence for EZH2, SUZ12 and H3K27me3 of cortical organoids across a developmental time course (from W2 to W10) from iPSC line 1. (E, F) Quantification of EZH2, SUZ12 and H3K27me3 intensity in nuclei (segmented based on DAPI) in the VZ and in CTIP2-positive N at W4 (E) and VZ and SVZ/CP at W6 (F) of cortical organoid culture relative to the VZ. Scale bars, 100 µm. Bar graphs represent mean values. Error bars represent SD; C, of 3 tissue samples from independent individuals; E, F, of 2–3 ventricles from at least 2 organoids. ** $p < 0.01$, * $p < 0.05$; Unpaired Student's *t*-test.

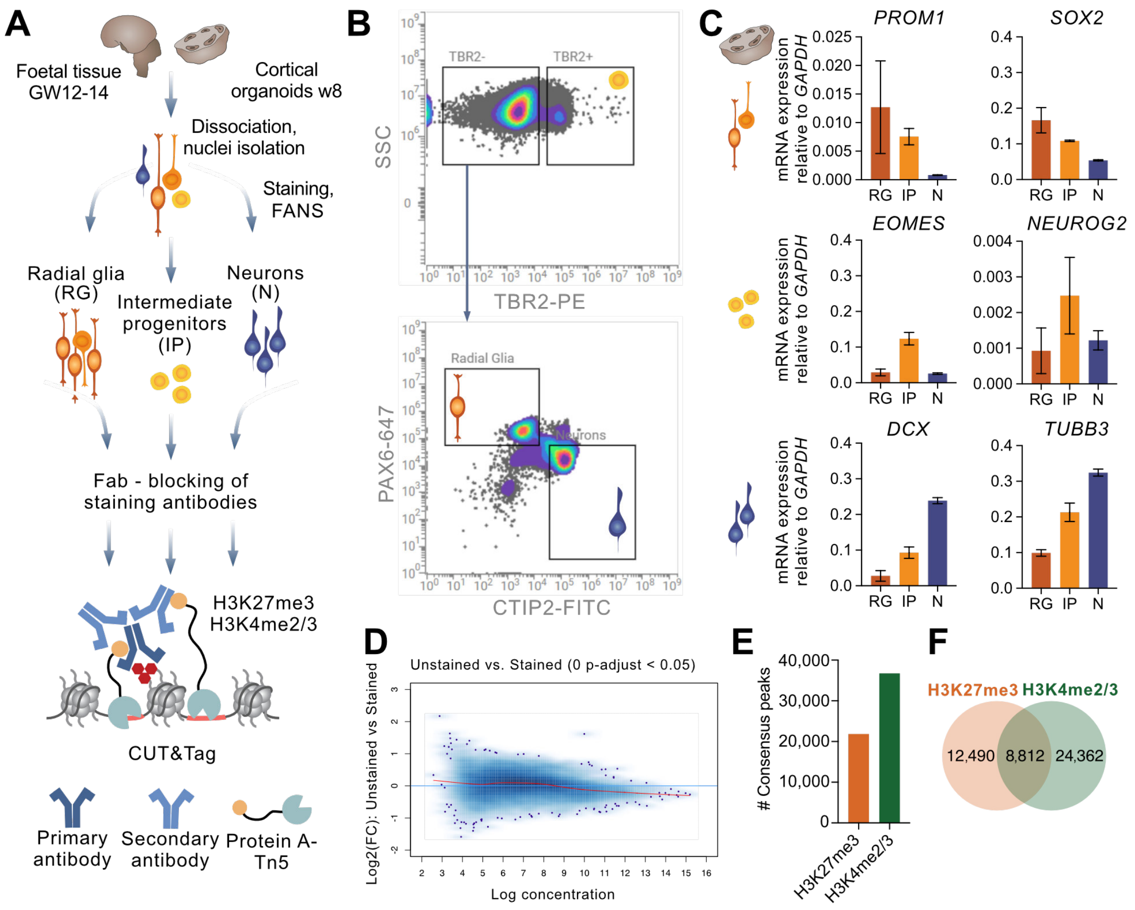


**Figure S4 related to Figure 3. Cell type-specific CUT&Tag on human foetal tissue and cortical organoids**.

(A) Experimental scheme for cell type-specific profiling of histone marks by CUT&Tag acquired for RG, IP and N from human foetal tissue (GW12–14; two tissue samples from independent individuals) and cortical organoids (W8). (B) Fluorescent activated nuclei sorting (FANS) plots depicting gating strategy of RG based on high levels of PAX6 and low levels of TBR2; IP based on high levels of TBR2, irrespective of other markers; and neurons based on high levels of CTIP2 and low levels of TBR2. Representative plots from human cortical organoids are shown. (C) Confirmation of cell type identity by mRNA expression analysis of marker genes characteristic of RG (*PROM1*, *SOX2*), IP (*EOMES*, *NEUROG2*) and neurons (*DCX*, *TUBB3*) for human cortical organoids. Bar graphs represent mean values. Error bars represent SD of three technical replicates. (D) MA plot illustrating differential enrichment of H3K27me3 between stained nuclei and unstained control nuclei, determined by DESeq2. Differential analysis did not identify any regions significantly enriched for H3K27me3 in the stained populations (p-adjust < 0.05, logFC > 1), indicating that staining of nuclei with sorting antibodies does not result in unspecific CUT&Tag enrichment. (E) Number of H3K27me3 and H3K4me2/3 consensus peaks generated for each histone mark individually, combining binding sites from all analysed cell types. Depicted are mean values from two independent biological replicates from human foetal tissue. (F) Venn diagram illustrating the overlap of H3K27me3 and H3K4me2/3 consensus peaks generated for each histone mark individually combining binding sites from all analysed cell types. Depicted are mean values from two independent biological replicates from human foetal tissue.

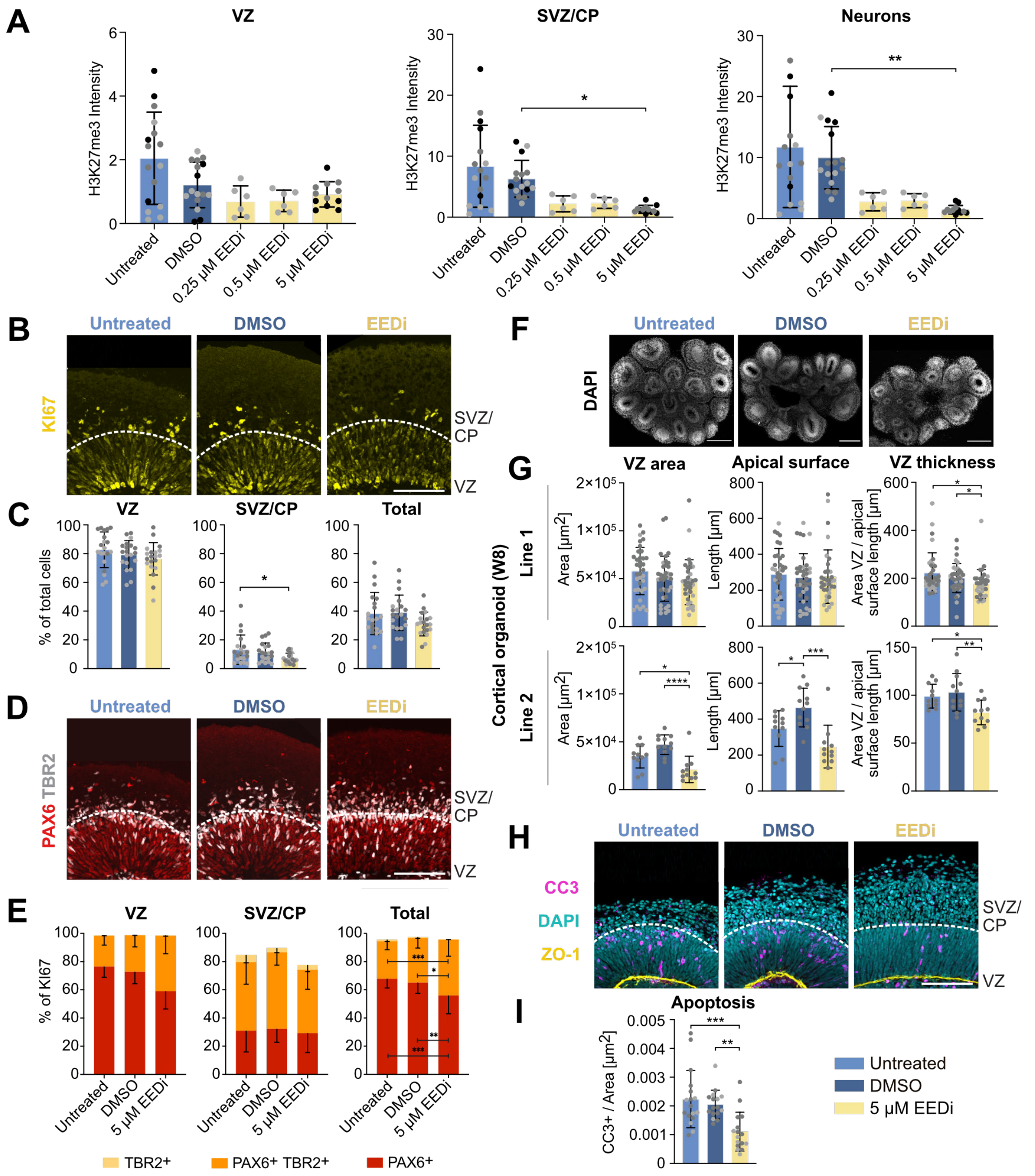


**Figure S5 related to Figure 4. Inhibition of PRC2 induces cell fate changes in human cortical organoids.**

(A) Quantification of H3K27me3 staining intensity in nuclei (segmented using DAPI) of VZ, SVZ/CP and CTIP2-positive neurons of cortical organoids (W8). Untreated and DMSO treated organoids serve as control. Treatment with 0.25 µM, 0.5 µM and 5 µM EEDi was applied for 5 weeks (W3 to W8). (B) Immunofluorescence of KI67 in cortical organoids (W8) from control (untreated, DMSO) and EEDi (5 µM) treatment conditions. (C) Quantification of KI67 as percentage of total cells (determined by DAPI) in VZ, SVZ/CP and total area. (D) Immunofluorescence of PAX6 and TBR2. (E) Quantification of PAX6 and TBR2 as percentage of total KI67+ cells in VZ, SVZ/CP and total area. (F) DAPI staining of whole cortical organoids. (G) Quantification of the VZ area (based on DAPI including the pseudostratified areas of ventricle-like structures facing the outside of the organoids), apical surface length (based on ZO-1 signal) and VZ thickness (calculated as the ratio of VZ area and apical surface length). Quantification was performed for organoids from two independent iPSC lines. (H) DAPI staining and immunofluorescence of the apoptosis marker CC3 and apical surface marker ZO-1. (I) Quantification of CC3 per total area. Scale bars, 100 µm (B, D, H) and 400 µm (F). Bar graphs represent mean values. Error bars represent SD; A, of 6–16 ventricles from at least 3 organoids; C, E, of 20 ventricles from at least 12 organoids; G, of 18–20 ventricles from at least 10 organoids (iPSC line 1) and 11–12 ventricles from at least 3 organoids (iPSC line 2); I, of 15–16 ventricles from at least 9 organoids from 2 organoid batches (indicated by different colours). ** $p < 0.01$, * $p < 0.05$; One-way (B, C, E, F) or two-way (D) ANOVA with Tukey's post-hoc test, (G) Kruskall-Wallis test with Dunn's post-hoc test.

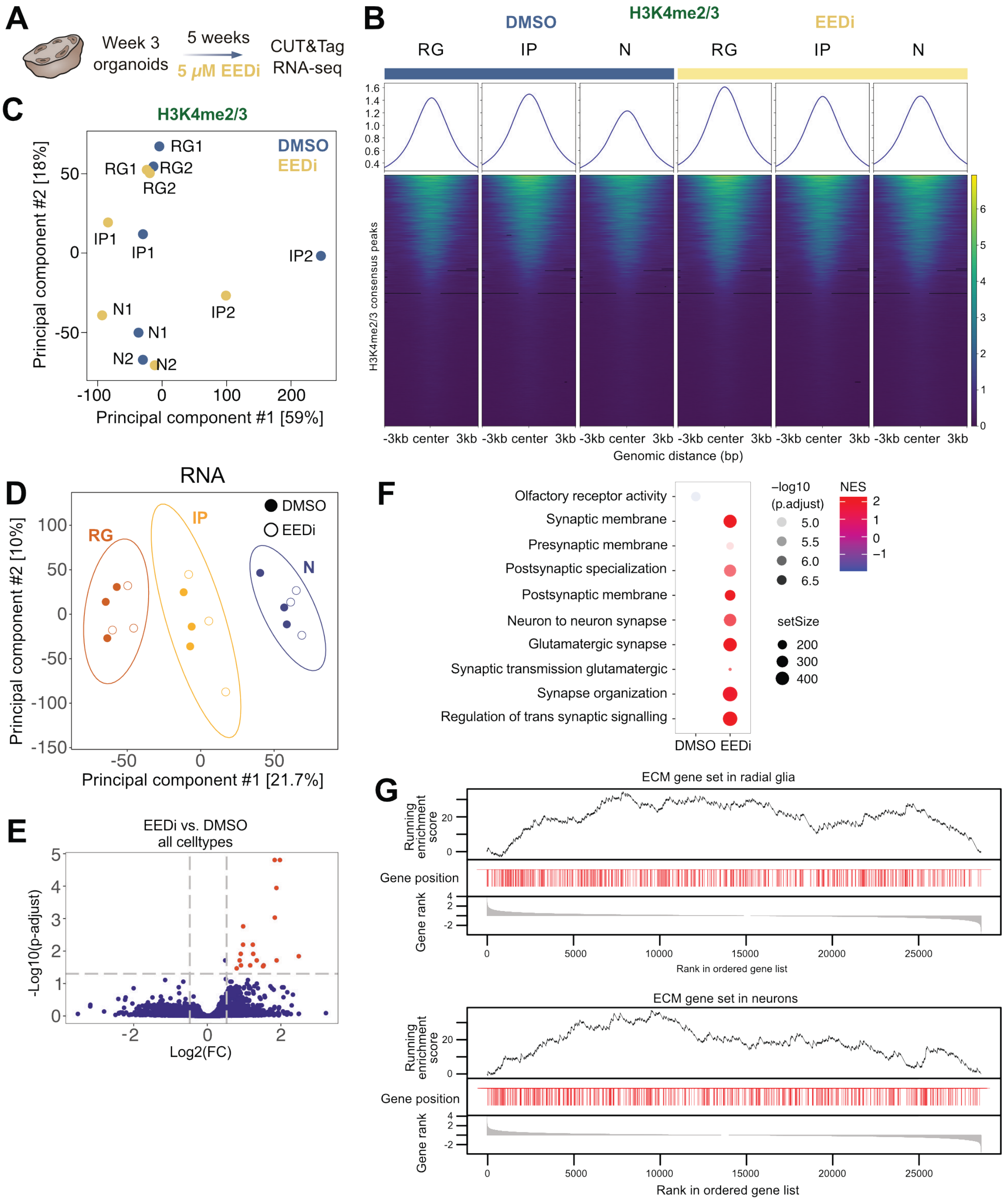


**Figure S6 related to Figure 6. Effect of PRC2 inhibition on H3K4me3 and gene expression.**

(A) Experimental scheme for pharmacological inhibition of PRC2 in human cortical organoids. (B) Profiles and heatmaps of the enrichment of H3K4me2/3 in RG, IP and N sorted from cortical organoids treated with EEDi or DMSO determined by CUT&Tag, in a set of consensus peaks. Each column represents the mean of two independent organoid batches. (C) PCA analysis for cell type-specific H3K4me2/3 acquired for RG, IP and N. Each dot presents a cell type from an independent organoid batch treated with EEDi or DMSO. (D) PCA analysis for cell type-specific gene expression analysed by RNA-seq for RG, IP and N. Each dot presents a cell type from three independent organoid batches treated with EEDi or DMSO. (E) Volcano plot displaying differentially expressed genes between EEDi and DMSO control condition across all cell types. Regions significantly enriched in one condition are marked in red (p-adjust < 0.05, logFC > 0.5). (F) GO term enrichment analysis for genes differentially expressed in EEDi compared to DMSO control. (G) Gene set enrichment analysis for ECM genes in RG (top) or N (bottom) in EEDi compared to DMSO control.

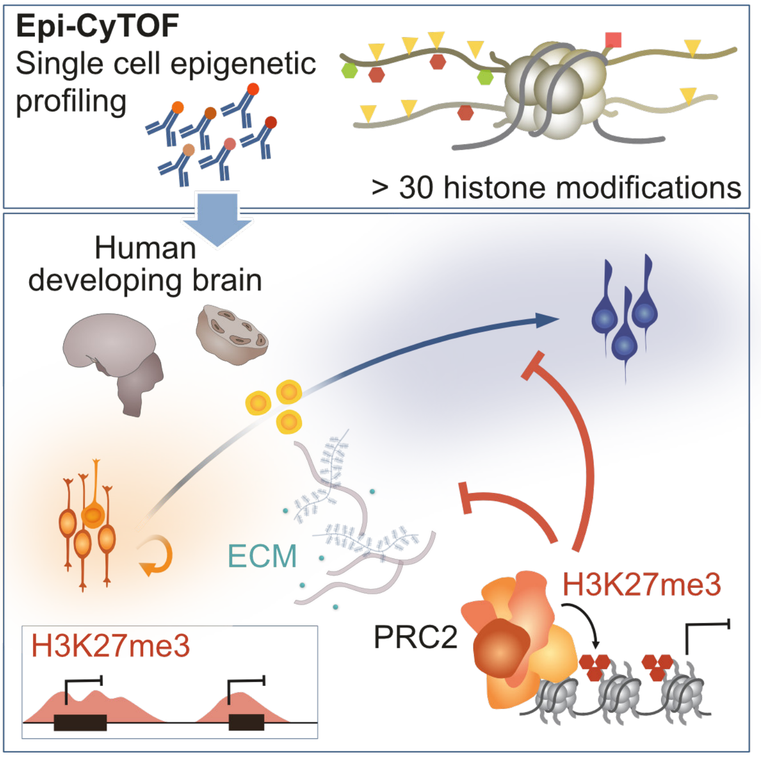


**Figure S7 related to Figure 6. Model of PRC2-mediated regulation of NPC fate in human corticogenesis.**

Schematic illustration of the Epi-CyTOF method allowing for single cell epigenome profiling of more than 30 histone marks, including H3K27me3. Subsequent cell type-specific profiling of the genome-wide H3K27me3 profiles not only identified neuronal differentiation, but also ECM genes as targets of PRC2. Inhibition of PRC2 in human cortical organoids resulted in a shift of NPC proliferation towards differentiation. Taken together, these findings suggest a dual role for PRC2 regulation in repressing neuronal differentiation. PRC2 simultaneously targets neurogenic genes cell-intrinsically, while also regulating the expression of ECM genes, thus contributing to the determination of neural cell fate in the human developing neocortex, by shaping the composition of the stem cell niche.
